# Mapping the dynamic transfer functions of epigenome editing

**DOI:** 10.1101/2021.01.05.425451

**Authors:** Jessica B. Lee, Leandra M. Caywood, Jennifer Y. Lo, Nicholas Levering, Albert J. Keung

## Abstract

Biological information can be encoded in the dynamics of signaling components which has been implicated in a broad range of physiological processes including stress response, oncogenesis, and stem cell differentiation. To study the complexity of information transfer across the eukaryotic promoter, we screened 119 dynamic conditions—modulating the frequency, intensity, and pulse width of light—regulating the binding of an epigenome editor to a fluorescent reporter. This system revealed highly tunable gene expression and filtering behaviors and provided the most comprehensive quantification to date of the maximum amount of information that can be reliably transferred across a promoter as ∼1.7 bits. Using a library of over 100 orthogonal epigenome editors, we further determined that chromatin state could be used to tune mutual information and expression levels, as well as completely alter the input-output transfer function of the promoter. This system unlocks the information-rich content of eukaryotic epigenome editing.

## Introduction

There is ample evidence that biological information can be encoded in the dynamics of signaling components and not just in their biochemical identities (Behar and Hoffmann, 2010; Cai et al., 2008; Dalal et al., 2014; Hansen and O’Shea, 2013; Hao et al., 2013; Imayoshi et al., 2013; Inoue et al., 2016; Purvis et al., 2012). Cells, with a limited number of components, utilize dynamic signal processing to perform sophisticated functions in response to complex environments. Transcription factors (TF) may be a particularly important archetype for this type of information transmission, as they are relatively low in diversity but must command many distinct and complex gene expression programs (Lee and Young, 2013). Indeed, through chemical and optogenetic approaches, the dynamics of TF nuclear-cytoplasmic translocation has been shown to control gene expression levels and population noise (An-adirekkun et al., 2020; Chen et al., 2020; Hansen and O’Shea, 2013; Rademacher et al., 2017). There is also evidence different promoters can transduce dynamic TF input signals into distinct output responses (Chen et al.; Hansen and O’Shea, 2016; Harton et al., 2019). Thus, developing a quantitative understanding of how dynamic TF signals are ultimately interpreted and processed by individual genes and promoters is clearly important.

Compelling analogies can be drawn: to information theory, with promoters analogous to information transfer channels; and to process control, with promoters acting as unit processes with dynamic input-output transfer functions. The nature of these channels or transfer functions might even be tunable by parameters such as promoter sequence, chromatin state, or three-dimensional chromatin topology. However, developing this type of robust quantitative framework poses considerable challenges. Mapping the transfer function of a single promoter seems ostensibly simple but faces the inherent technical difficulties of controlling dynamic properties of biological systems. The complex diversity of eukaryotic chromatin presents yet another formidable barrier. More specifically, there are three particularly pressing challenges. First, there is a broad range of dynamic input and output parameters that is technically challenging to access, control, and characterize. Second, as each individual promoter can be regulated by multiple distinct TFs and chromatin regulators (CRs), pleiotropic effects can confound global perturbations to nuclear TF levels or chromatin state. Finally, there are hundreds of distinct CRs that can alter how promoters interpret TF signals, resulting in a large experimental space to explore (Kouzarides, 2007; Li et al., 2007).

To address these challenges we engineered both dynamic and static epigenome editors that bypass pleiotropic issues due to their locus specificity and thereby provide insight into the causal impacts of CRs and TFs on transcription (Bintu et al., 2016; Keung et al., 2014; Park et al., 2019; Polstein and Gersbach, 2015). To study the effects of TF signal dynamics on transcription, we employed an optogenetic system that dynamically recruited the transactivator VP16 to a genomically-integrated reporter. By pairing the optogenetic system with programmable Arduino-controlled LED arrays and single-cell fluorescence measurements by flow cytometry, we were able to efficiently capture and screen a large parameter space of dynamic inputs.

Using this experimental platform, we comprehensively mapped transcriptional outputs in response to 119 different optogenetic inputs that modulated the amplitude, frequency, and pulse width of VP16 recruitment. Input conditions with the same total signal but different dynamic parameters yielded outputs with over an order of magnitude difference and, therefore, acted as a filter. A kinetic model was developed to describe the complex transfer function captured by the experimental data, including the filtering behavior. To further understand the reliability of the information transfer, we applied information theory to the single cell distribution data and estimated the maximum amount of information transmittable through each input mode—as well as with all input modes combined—with frequency modulation carrying the greatest amount of transmittable information and amplitude the least. Finally, we asked if co-recruitment of CRs to the promoter could alter its transfer function without any alteration to the promoter sequence. 101 CRs were constitutively recruited to the promoter. Many of them altered the transcriptional response to dynamic VP16 inputs, including exhibiting new complex types of transfer functions such as band-pass, low-pass, and high-pass frequency filtering. In addition, co-recruiting CRs with VP16 tuned the maximum amount of information that was transmittable through the single promoter. This study reveals the information-rich nature of eukaryotic transcription even at just a single promoter, implicates an important interplay between dynamics and chromatin, and also provides quantitative synthetic biology, modeling, and information theory frameworks to understand and predict complex transcriptional transfer functions.

## Results

### Optogenetics provides complete access to the dynamic parameter space (Figure 1 and S1)

We developed an optogenetic system to recruit epigenome editors to a synthetic transcriptional reporter in arbitrary dynamic patterns (Figure 1A). A *CYC1* promoter drove expression of an mCherry reporter and was integrated into the *LEU2* locus of *Saccharomyces cerevisiae*. The *CYC1* promoter contained two identical binding sites (GAGTGAGGA) recognized by an engineered zinc finger (ZF) array “ZF43-8” and an orthogonal binding site recognized by ZF array “ZF97-4” (TTATGGGAG) (Keung et al., 2014; Khalil et al., 2012). In addition, we fused ZF43-8 to cryptochrome 2 (*ZF-CRY2*) and cryptochrome-interacting basic helix-loop-helix to the transcriptional activator *VP16* (*CIB1-VP16*) and placed their expression under ATC and IPTG control (Keung et al., 2014), respectively. CRY2 binds CIB1 when exposed to blue light and dissociates upon light removal (Kennedy et al., 2010; Liu et al., 2008). This system has high temporal resolution with an association half-life of seconds and dissociation half-life of ∼5 minutes (Rademacher et al., 2017). We also tested other optogenetic systems, different N-C terminal fusions, and several induction drug concentrations (see Figure S1). The final system was chosen for its robust activation with light and minimal activation without light. To deliver the light signals, an Arduino Due controlled individually addressable blue LEDs (wavelength= 455-465 nm) in a 96-well format.

**Figure 1.**
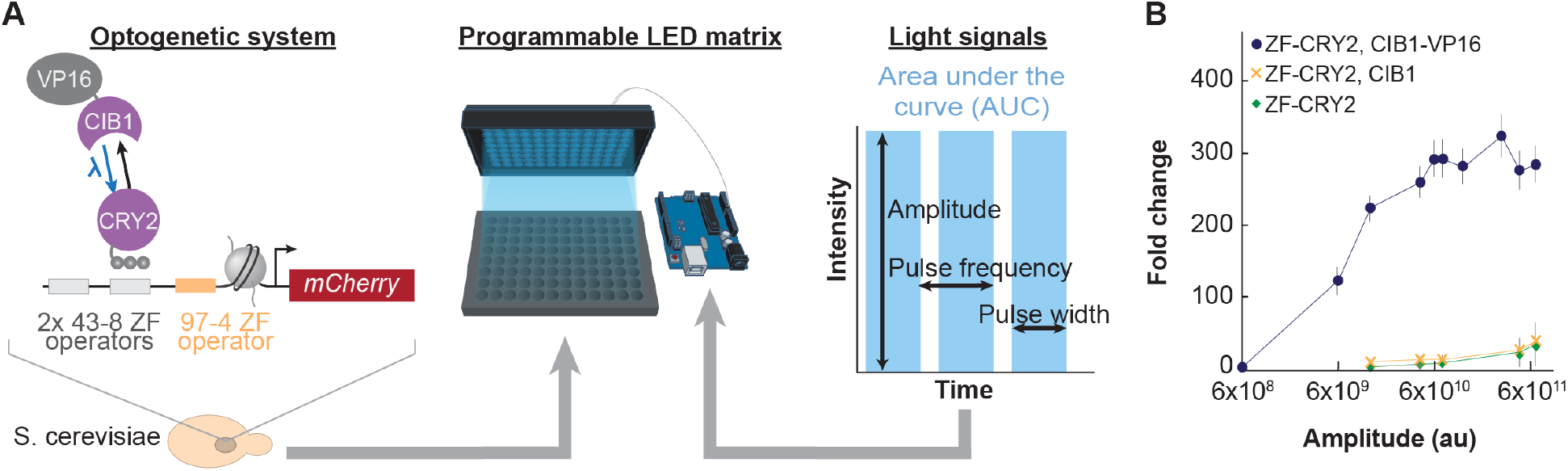
Optogenetics provides complete access to the dynamic parameter space. (A) Schematic of genetic and hardware systems. The optogenetic system was expressed in *S. cerevisiae* (left). ZF-CRY2 targeted operators placed upstream of a minimal *CYC1* promoter driving the expression of mCherry. In the presence of blue light, CIB1-VP16 binds ZF-CRY2 and disassociates without blue light. Parameters of amplitude, frequency, and pulse width (right) were varied using a custom Arduino-controlled, individually addressable LED matrix (center). The area under the curve is defined as the (amplitude) x (frequency) x (pulse width) x (duration of experiment). (B) Fold change in fluorescence for various light intensity amplitudes for a constant, 14-hour light pulse. Fold changes for control strains, ZF-CRY2 and ZF-CRY2+CIB1, are also shown. Dots represent mean, error bars are standard error of the mean (s.e.m.) for three biological replicates.

To accurately map the effects of different dynamic input light patterns on eukaryotic transcription, the system must operate at sub-saturation. Therefore, we first determined the dynamic range of the system and identified sub-saturation light amplitudes (i.e. intensities). We exposed the cells to a range of light pulse amplitudes that were constantly on for 14 hours. The resultant fold change was measured by flow cytometry and was defined as the median fluorescence of cells incubated with a given light pattern divided by the median fluorescence of cells incubated without light (Figure 1B). The output signal saturated at intensities above 6×10^10^ au. Importantly, control cells expressing only ZF-CRY2 or ZF-CRY2 with CIB1 exhibited much lower levels of activation, especially at sub-saturating levels. For further studies modulating all three dynamic parameters (amplitude, frequency, and pulse width), we chose sub-saturation amplitudes below 6×10^10^ au to ensure both comprehensive coverage of the dynamic parameter space and to minimize any activation due to ZF-CRY2 alone.

### 119 dynamic signals provide comprehensive map of a eukaryotic transfer function (Figure 2 and 2S)

Frequency, amplitude, and pulse width modulation (F, A, and PW, respectively) present a large combinatorial space which is challenging to capture experimentally; yet, it is crucial to do so in order to understand the transfer functions of eukaryotic promoters and to generate quantitative and predictive models. With the programmable LED array paired with flow cytometry, we delivered 119 distinct input signal patterns to yeast cultured in 96-well format and measured mCherry reporter end-point fluorescence after 14 hours. These patterns included 5 sets of conditions with PW held constant at 5, 120, 600, 1800, or 3600 seconds. We chose this range of PWs to include timescales similar to those found for several pulsatile TFs in *S. cerevisiae* (Dalal et al., 2014). For each PW, four amplitudes (6×10^9^, 1.2×10^10^, 4×10^10^, and 6×10^10^ au) and 5-6 frequencies (between 2×10^−5^ and 1×10^−1^ sec^-1^) were delivered to the cells. We defined the total input signal, or area under the curve (AUC), as the product of F, A, and PW for the duration of the experiment. The throughput of the system allowed us to measure thousands of cells as well as four biological replicates per condition.

**Figure 2.**
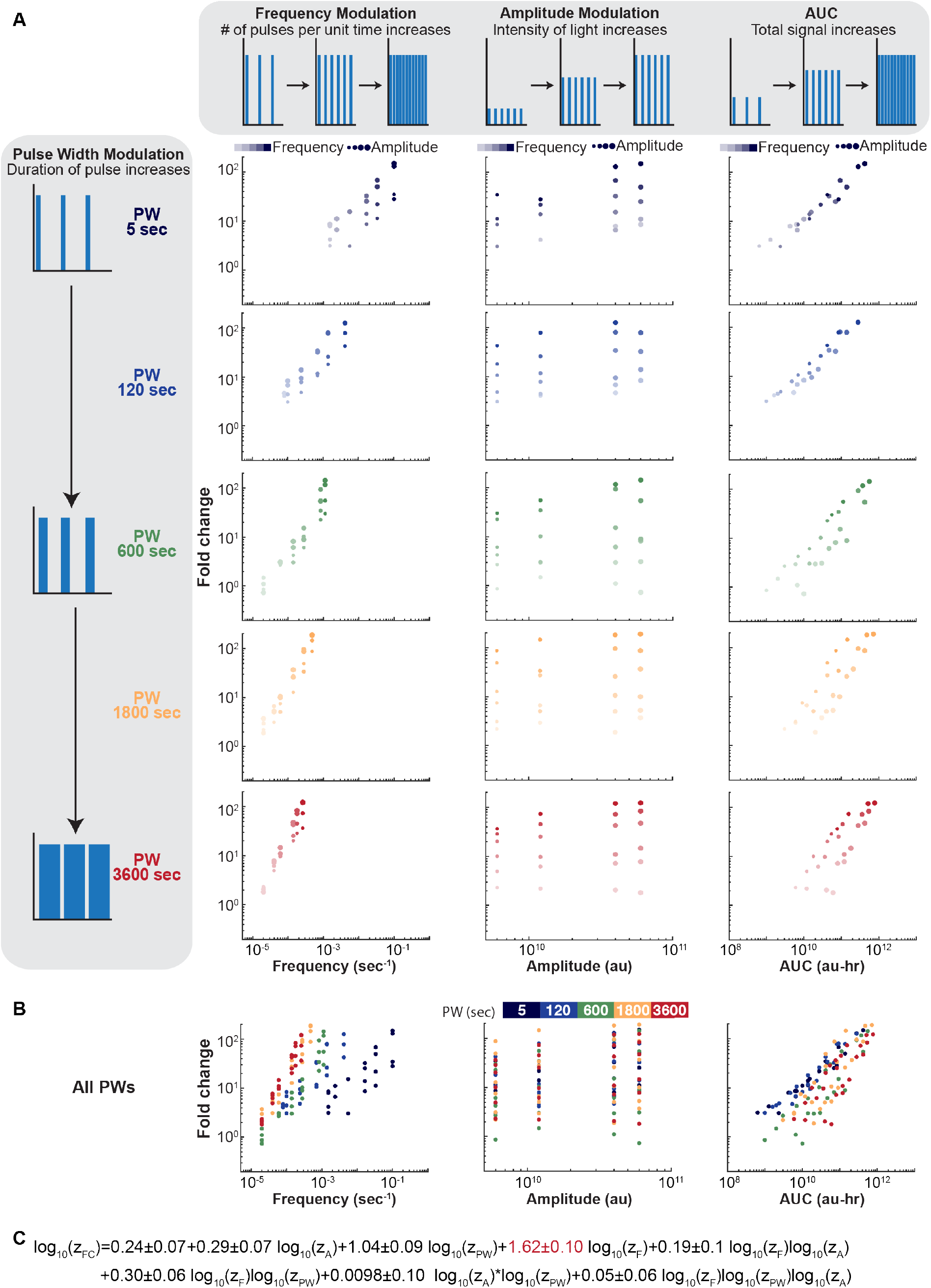
119 dynamic signals provide comprehensive map of a eukaryotic transfer function. (A) Schematics illustrating each mode of modulation (PW, F, and A) and AUC are shown along the top or left. Slope of fold change vs F increases with PW. Fold change exhibits only slight increases with A when PW and F are held constant. Fold change response collapses onto a line when plotted vs AUC for 5 and 120 sec PWs but not for higher PWs. (B) All conditions are plotted together, further emphasizing the filtering behavior at constant AUCs (right). Fold change=(median fluorescence with blue light condition-autofluorescence)/(median fluorescence without blue light-autofluorescence). Each dot is the mean of 4-8 replicates. (C) Fitted linear regression equation using the z-transformed parameters (z_A_, z_PW_, z_F_). The coefficient for z_F_ is significantly higher than the others (p=0.05).

As expected, mCherry expression increased with A, F, PW, and AUC (Figure 2A-B). However, while F and PW had strong effects on mCherry output (Figure 2A, left column), A had much weaker effects at both low and high regimes of mCherry output (middle column). To quantify the relative effect of each light parameter on mCherry expression, we first standardized the A, F, PW, and resulting mCherry fold change using z-transformation. This allowed comparison between the coefficients of each input mode within a regression model (Schielzeth, 2010). We then fit the standardized data to a linear regression model, given in Figure 2C (R^2^=0.92, p=5.29e-57). This linear model confirmed that frequency had the largest coefficient and therefore greatest effect on fold change, while amplitude had the weakest effect.

The fact that the system did not respond equally to each mode of modulation suggested that the system might exhibit filtering behaviors, where input signal patterns that share identical input AUCs but through different weightings of A, F, and PW, could yield distinct output levels. Indeed, this signaling filtering property was observed over a wide range of AUCs (Figure 2A and 2B, right column). This indicates that this system has inherent signal-filtering capabilities because F, PW, and A do not have equally proportional effects on mCherry expression. We found that filtering was not an artifact of measuring mCherry fluorescence at different time points after the last light pulse was delivered; the same filtering was observed even when the timing of the last light pulse was shifted relative to the end of the experiment (Figure S2).

### Model captures system behavior and filtering (Figures 3 and S3)

To our knowledge, our experimental system provided the largest and most comprehensive set of dynamic data to date. We therefore asked if it could inform the development and architecture of mass action models of eukaryotic transcription, and if this model could subsequently provide additional insights into the filtering property of the system. We tested several previous models (Benzinger and Khammash, 2018; Chen et al., 2020; Hansen and O’Shea, 2013; Harton et al., 2019) along with some modifications and found that a four state model (probability of each state is represented by P_unbound_, P_bound_, P_inactive_, and P_active_) best fit the experimental data with R^2^=0.835 (Figure 3, Figure S3). Compared to a similar prior model that differed in the number of Hill functions incorporated into the rate constants (Hansen and O’Shea, 2013), this model exhibited less intense fluorescence oscillations over time, producing a ‘smoother’ response (Figure S3B). Because we did not see large differences when light pulses were shifted to all start (Figure 2) or end (Figure S2) at the same time, we determined that the actual fold change did not exhibit large oscillations and was better represented by this four-state model. We did explore incorporating various numbers of Hill functions into the model as was done in other models of eukaryotic transcription (Figure 3C). We found that the inclusion of a single Hill function yielded the best fit.

**Figure 3.**
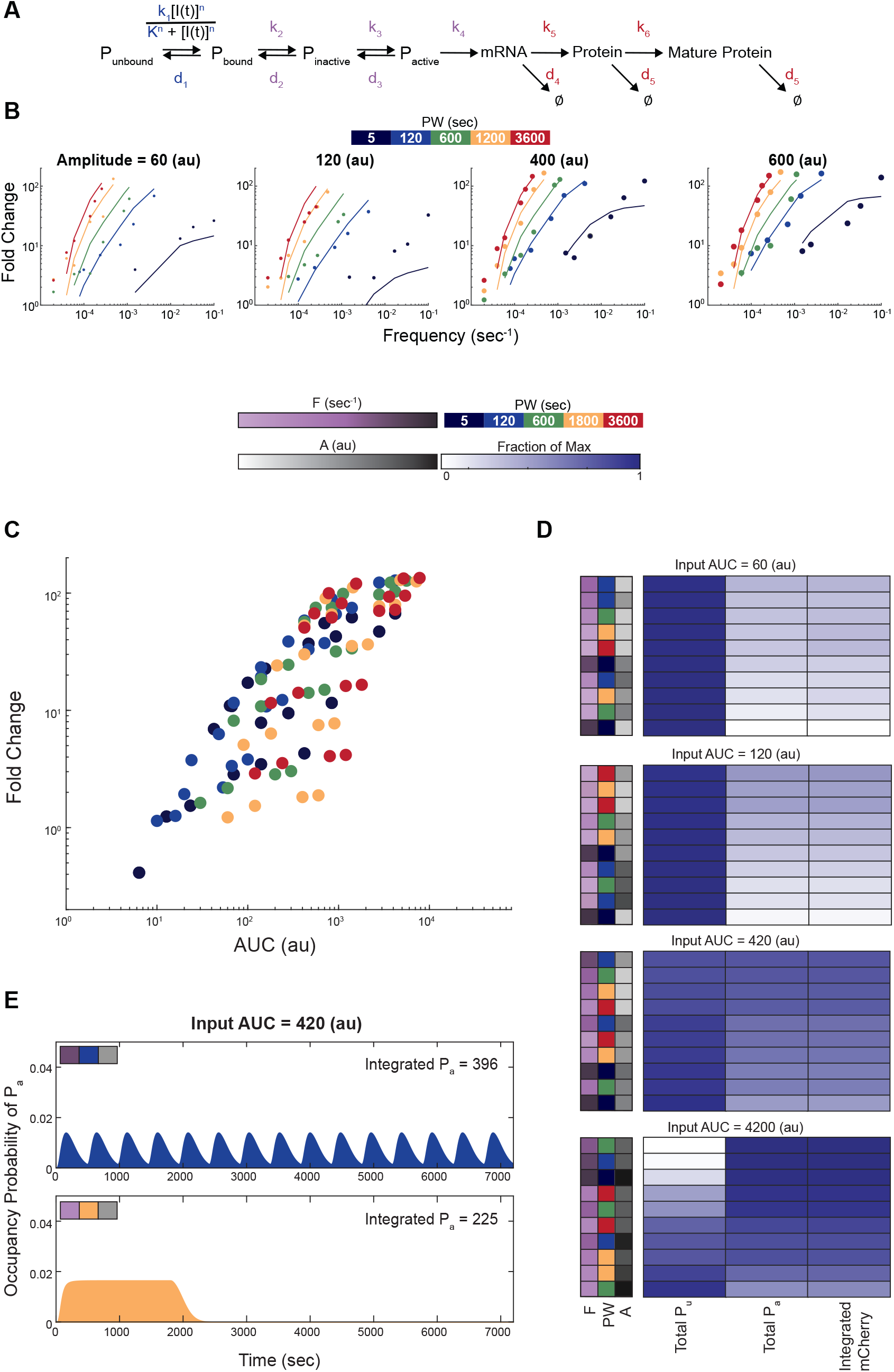
Model captures system behavior and filtering. (A) Schematic of the four-state model used to fit the experimental data. Blue parameters (k_1_, d_1_, K, and n) were adapted from Rademacher et al. Red parameters (k_5_, d_4_, k_4_, d_5_, and k_6_) were adapted from Hansen and O’Shea. Purple parameters (k_2_, d_2_, k_3_, d_3_, and k_4_) were varied and fit to experimental data. P_unbound_, P_bound_, P_inactive_, and P_active_ represent the probabilities of each promoter state. (B) The resulting fold changes for the model using the best-fitting parameter set (R^2^=0.835) are shown as lines. Experimental data are dots. (C) Fold change generated by the model shows close similarity to the filtering observed in the experiment in Figure 2B. (D) Heat maps show total integrated occupancy of P_unbound_, P_active_, and total integrated mCherry with the input light pattern indicated on the left, for four different AUCs. Values were log transformed and normalized to the max value within all heat maps. (E) Holding A and AUC constant, the integrated occupancy of P_active_ is higher for higher F conditions.

The filtering behavior observed in the experimental data, was an interesting feature of our optogenetic system. We therefore asked if our four-state model also captured this behavior. Indeed, our model reflected this filtering behavior closely (Figure 3C). Interestingly, the model suggested that filtering may arise from a combination of different mechanisms, as a variety of distinct input patterns with the same AUC gave rise to distinct outputs (Figure 3D). One contributor may be that the decay of promoter occupancy is not immediate, and so at high F there is an accumulation of ‘extra’ promoter occupancy (Figure 3E).

### Single cell measurements capture total population noise (Figure 4)

While understanding and mapping the transfer functions of a system is important, the reliability in achieving the same output repeatedly over time or within a population of cells in response to the same input signal is equally important. This reliability is characterized by the noisiness of gene expression. Gene expression noise is an important and inherently stochastic process due to the low copy number of genes (Eldar and Elowitz, 2010; Elowitz and Leibler, 2000; Maheshri and O’Shea, 2007) and, together with cell-to-cell variability in general cellular components, creates a distribution of single-cell outputs for each unique input (Elowitz and Leibler, 2000; Grabowski et al., 2019; Gregor et al., 2007; Rosenfeld et al., 2005; Tkacik et al., 2009). As noise plays an important role in determining the reliability of a system and the fidelity of transmitted information, we first quantified how noise in our system was affected by F, A, and PW modulation. We calculated the robust coefficient of variation (CV) of the population for all 119 input light conditions (Figure 4). Of note, the CV was much lower and nearly constant for all AUCs and Fs when mCherry fluorescence was normalized by size using FSC-A (Figure S4). This phenomenon agrees with prior work showing cell size as a major contributor to noise (Bar-Even et al., 2006; Newman et al., 2006), with removal of this noise due to size also providing an estimation of intrinsic noise (Newman et al., 2006).

**Figure 4.**
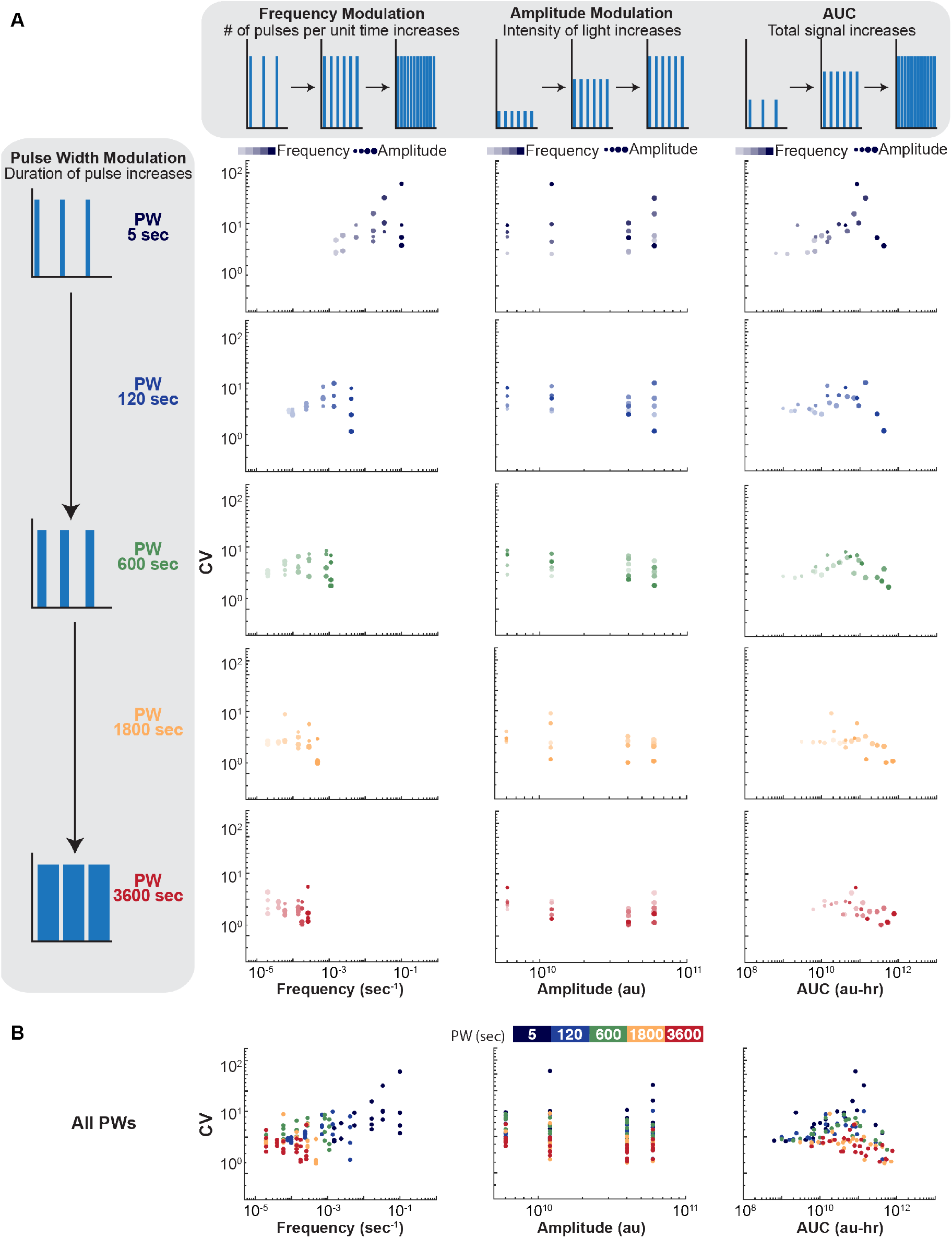
Single cell measurements capture total population noise. (A) The robust coefficient of variation (CV) calculated for 119 input light conditions. Schematics illustrating each mode of modulation and AUC are shown along the top or left side. Dot color, size, and shade correspond to PW, A, and F, respectively. (B) All PWs are plotted together. Each dot is the mean of 4-8 biological replicates.

### Quantifying the contribution of signaling dynamics to maximum mutual information (Figure 5)

With an understanding of the noise within our system, we next used the fluorescence distributions to determine the reliability of information transfer. Borrowing concepts from information theory, the capacity or reliability of an information transmission system can be quantified as the maximal mutual information (MI) (Shannon, 1948). In previous studies, MI has been used to quantify the signaling fidelity in terms of the maximum number of input values a cell can accurately resolve in the presence of noise (Cheong et al., 2011). Systems with MI less than 1 bit have a significant amount of overlap in their gene expression output distributions and can only resolve two inputs. In contrast, systems that can resolve multiple inputs have minimal overlap in the output distributions and have MI greater than 1 bit (Figure 5A). Recently, Hansen and O’Shea used MI to characterize the response to Msn2p signaling using F or A modulation. They observed that a single promoter could reliably distinguish three states (∼1.58 bits) (Hansen and O’Shea, 2015). It is unclear if this was the maximum possible for a single promoter, and whether combining modes of modulation could provide a higher estimate of information limits.

**Figure 5.**
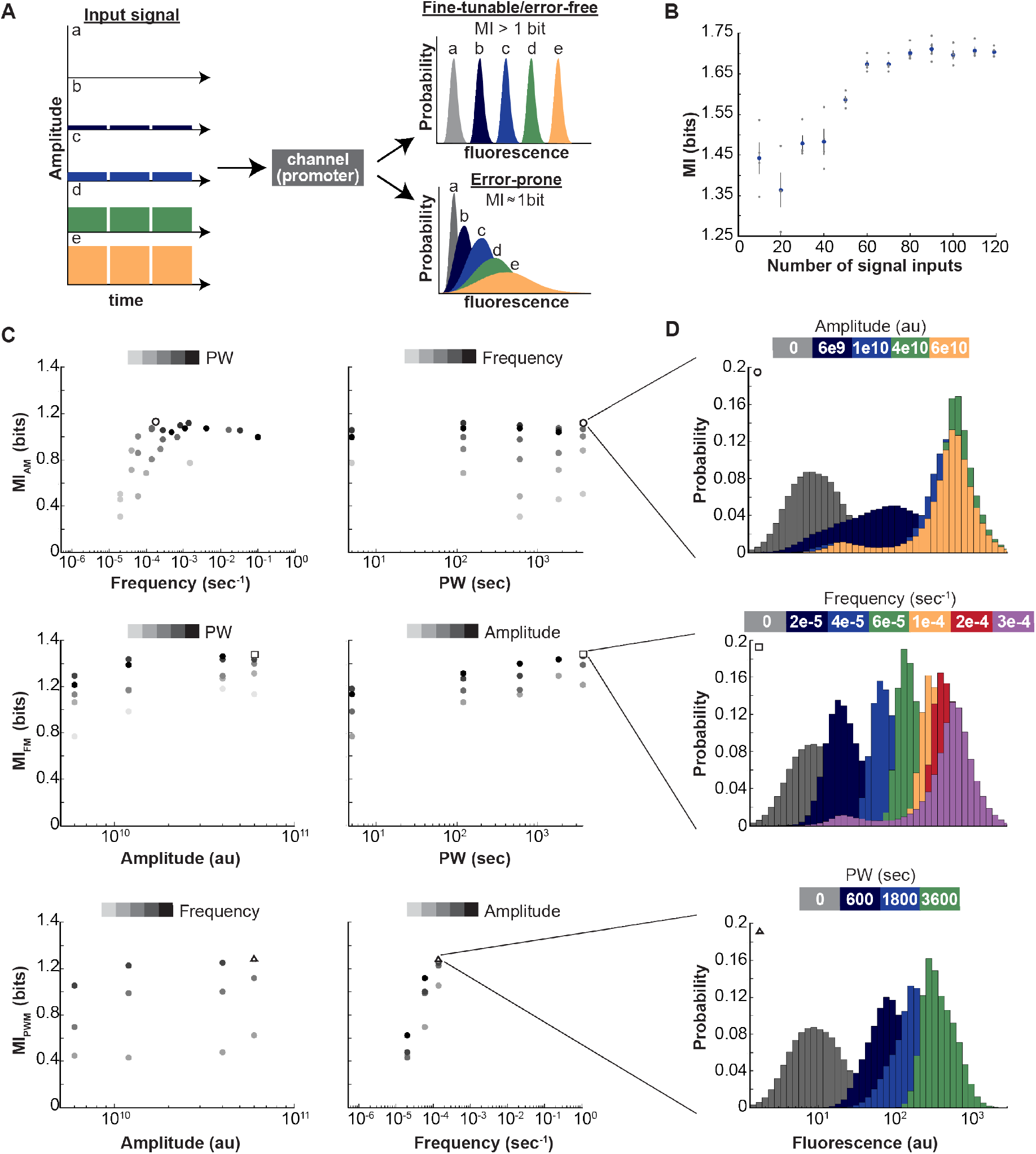
Quantifying the contribution of signaling dynamics to maximum mutual information. (A) Schematic showing potential signal inputs (left) for a promoter that is modelled as a noisy channel and two extreme cases of possible outcomes (right). With distinct outputs, the maximal mutual information (MI) is greater than 1 bit and the system is error-free and fine-tunable. With outputs with a large amount of overlap, the MI is less than or equal to 1, and the system is error-prone. (B) Plot of MI for all three modes of modulation as the number of signal inputs increases. The total fold change range was held constant for all combinations of inputs. Error bars are s.e.m., n=4 biological replicates. (C) MI for the different modes of modulation. Each dot is the mean of 4 biological replicates. (D) Fluorescence histograms of single cell distributions for different modes of modulation for the parameter set resulting in the highest MI for the specified mode, indicated by O, □, or Δ in (C) and (D).

As we have mapped a large number of input-output responses for all three dynamic modes of modulation (F, A, and PW), we asked what the information capacity limit of our single promoter was. To do this, we randomly selected subsets of the 119 different input signaling patterns and calculated the MI. Importantly, the same overall dynamic range of mCherry expression was maintained for all subsets of signal input patterns. We repeated this process for increasing numbers of input signaling patterns per subset and found that the MI started around 1.45 bits and increased before plateauing near an MI of 1.7 bits (Figure 5B). This indicated that the MI was dependent on the number of inputs and required a large screen of the parameter space to measure.

Biological signaling pathways can encode information through the A, F, or PW of a shared signaling molecule (Batchelor et al., 2011; Hao and O’Shea, 2012; Purvis and Lahav, 2013). However, it is unclear which method is the most reliable. Hansen and O’Shea showed that promoters that bind Msn2p have higher information transduction capacities using A rather than F modulation (Hansen and O’Shea, 2015). However, we found that the MI for each mode of modulation (M) depended on the constant values of the other two parameters (Figure 5C). For example, the MI_AM_ increased at low F and leveled off at a F dependent upon the PW (Figure 5C, top-left). As the PW increased, the F with the highest MI_AM_ decreased. For F modulation, there was a less pronounced increase in MI_FM_ as A or PW increased. MI_PWM_ showed little increase with A but a large increase with F.

When comparing the maximum MI for each mode of modulation, AM was less reliable (1.12 bits) than PWM (1.23 bits) and FM (1.48 bits). This agrees with our previous assessment that F had the greatest effect on fold change. The histograms of the outputs provide some insight into why AM had relatively low MI (Figure 5D). For AM, the amplitudes of 1.2×10^10^, 4×10^10^, and 6×10^10^ au had a high degree of overlap and were near saturation when the PW and F were at high values. Additionally, the 6×10^9^ au histogram had a very broad peak (CV=0.57± 0.02, s.e.m.), which decreased the MI. In fact, low A conditions exhibited broad distributions in general. In contrast, both PWM and FM outputs were more distinguishable, even when the other parameters were at their maximum values. They also exhibited tighter distributions, and therefore had higher MIs. The differences between our system and Msn2p of Hansen and O’Shea could be attributed to several factors, including: 1) different mechanisms of activation of Msn2p and VP16; 2) different promoter sequence structure, e.g. location of binding sites relative to transcription binding site; 3) differences in the binding kinetics; and 4) different genomic location of the reporter and therefore different initial chromatin state of the promoter.

### Chromatin regulators tune maximum information content

We were able to map the transfer function and quantify the reliability of information transfer for our single synthetic promoter. However, prior work has shown that distinct promoters can exhibit different regulatory behaviors including distinct dynamic ranges of expression, activation kinetics, and noise (Hansen and O’Shea, 2013, 2015; Hao et al., 2013), with a likely explanation for these differences being distinct chromatin states. Yet, an individual promoter can also exist in diverse chromatin states that might alter the way it responds to input signals (Hansen and O’Shea, 2013; Li et al., 2007). We hypothesized that chromatin state, defined by a complex combination of features including nucleosome positioning, nucleosome modifications, three-dimensional topology, and the presence of diverse chromatin regulating proteins could alter both the maximum information transmittable by a single promoter and the nature of its transfer function without any change to its DNA sequence.

To determine if the chromatin landscape could change the MI without altering the promoter sequence, we created a library comprised of ZF97-4 fused to each of 101 chromatin regulators (CRs) chosen for a diversity of putative activities and membership in a variety of protein complexes, e.g. SAGA, TFIID, SWI/SNF. These CRs were constitutively recruited to the *CYC1* promoter (Figure 6A). VP16 was then dynamically recruited. Given the large number of yeast strains in this library, we focused on varying F while keeping A and PW constant, as frequency had yielded the greatest MI when VP16 was recruited alone. Four input Fs were measured: 0 (i.e. dark), 6.7×10^−4^, 3.3×10^−2^, and 1×10^−1^ sec^-1^. A and PW were held constant at 6×10^10^ au and 5 sec, respectively.

**Figure 6.**
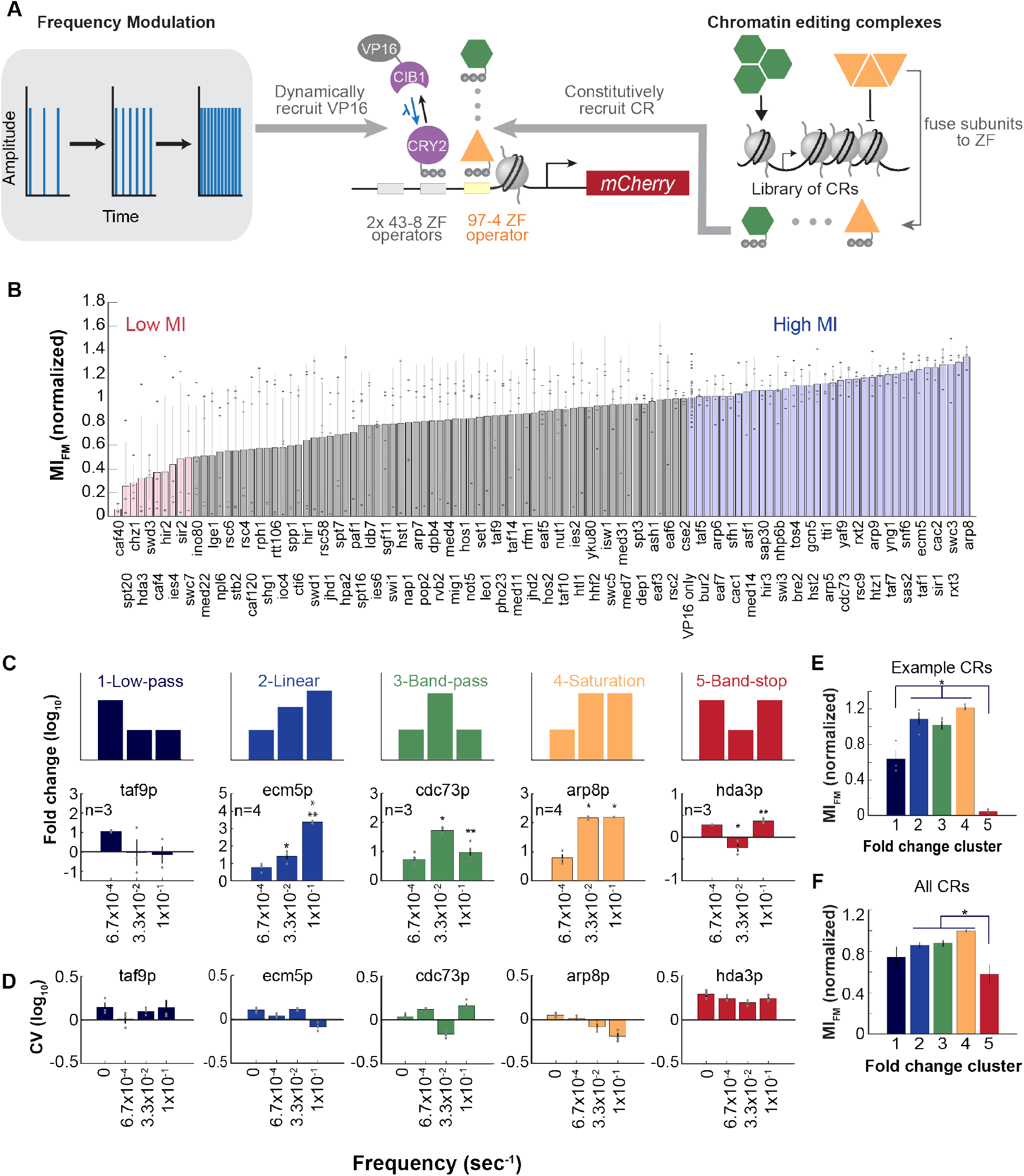
Chromatin regulators tune MI and morph signal filtering transfer functions achievable by a single promoter. (A) Eukaryotes utilize a diverse set of protein complexes capable of editing the epigenome. 101 subunits of these complexes were each fused to a 97-4 zinc finger (A, right). This allowed recruitment of the chromatin regulators to the same promoter (center) as the optogenetically-controlled VP16 (left). (B) Maximal mutual information for frequency modulation using frequencies 0, 6.7×10^−4^, 3.3×10^−2^, and 1×10^−1^ sec^-1^ for the library of ZF97-4-CRs. Error-bars are s.e.m. Gray dots are biological replicates. N=3-4. VP16 only strain is 48 technical replicates. (C) The strains containing ZF97-4 CRs were grouped into five clusters (C, top). Each exhibited a different type of signal filtering. Example CRs for each cluster (C, bottom). *p<0.05 compared to frequency=6.7×10^−4^ sec^-1^, **p<0.05 compared to frequency=3.3×10^−2^ sec^-1^. (D) Noise for the example CRs for all frequencies, including 0 (e.g. dark). (E) Average MI_FM_ for example CRs shown in (C). *p<0.05, Tukey-Kramer post-hoc. (F) Average MI_FM_ of all strains within each cluster. *p<0.05, Tukey-Kramer post-hoc. Error bars are s.e.m.

With these experimental conditions, we obtained an MI_FM_ value for the promoter with each distinct CR recruited (Figure 6B). The MI_FM_ ranged from 0.064± 0.02 (s.e.m.) for Caf40p to 1.34± 0.04 for Arp8p. The values of MI_FM_ in Figure 6 are lower than those shown in Figure 5 because only four distinct input light conditions were tested for each CR instead of the 6-7 used in Figure 5. To provide a means for mutual comparison, each MI_FM_ was normalized to the MI_FM_ of the yeast strain without ZF97-4. Of note, there were strains with large MI_FM_ variability among the biological replicates (for example, Hir2p and Nap1p). This is not unexpected because the VP16-only strain exhibited relatively large variability when fewer inputs were used to calculate MI. As more inputs were included, the MI became more consistent between replicates (Figure 5B). However, even with only four distinct inputs, it was apparent that CRs affected the MI_FM_ (p=1.18×10^−7^, ANOVA). This suggests that chromatin may regulate how “fine-tunable” a gene is and how much information can be transmitted reliably via transcription. To gain further insight, we also clustered CRs that had low (less than 0.5) and high (above 1.15) MI_FM_. Through gene ontology, we found significant enrichment within the low MI_FM_ cluster of CRs involved in RNA catabolic process, mitochondrion organization, organelle fission, peroxisome organization and regulation of translation (Fisher exact test with Bonferroni correction, p<0.05). Although not statistically significant, CRs involved in DNA recombination and response to DNA damage had a the highest MI_FM_ values, suggesting these CRs may be recruited in natural situations to enhance the reliability of signals to induce DNA repair (See Figure S5).

### Chromatin regulators diversify the transfer functions achievable by a single promoter

Previous studies have shown that different promoters can exhibit distinct transfer functions (Hansen and O’Shea, 2013, 2016; Harton et al., 2019), not just alterations in MI. We asked if constitutive recruitment of CRs could alter the transfer function of a single promoter without changes to the DNA sequence. In particular, we asked whether CRs alter the qualitative signal filtering properties of the promoter. For example, can CRs allow the promoter to respond preferentially to low or high frequency input signals, and not just shift the dynamic range of the output response?

To address this question, we clustered all sets of biological replicates by their pattern of output responses to low, medium, and high frequency input signals. This was performed in an unsupervised manner using k-means clustering. Each cluster exhibited different filtering behaviors: low pass, linear, band-pass, saturation, and band-stop. To our knowledge, this is the first demonstration that the same promoter can exhibit multiple types of filtering, tunable by CRs. Example CRs for each cluster are shown in Figure 6C-F. It should be noted that only Hda3p had all of its biological replicates grouped into the band-stop cluster which may be a relatively rare filtering property. Furthermore, band-stop transfer functions (e.g. Hda3p) had significantly lower MI_FM_ than the other clusters (Figures 6 D-F); this is most likely due to the narrow dynamic range and high noise. The noise of each strain was generally inversely proportional to the fold change (Figure 6D) and therefore also exhibited filtering behavior. When assessing the MI for each cluster, the trend suggested that CRs may need to sacrifice MI and information transmission capacity to achieve more exotic signal filtering properties like low pass and band stop filtering (Figure 6E-F).

## Discussion

Many transcription factors exhibit pulsatile behavior in response to stress. We addressed the question of how an individual gene interprets this type of dynamic input signal. Using optogenetics to induce 119 distinct dynamic input signals, we comprehensively mapped the transfer function of an individual promoter as well as the associated noise and reliability of information transmission as a function of dynamic parameters. A four-state kinetic model was able to capture this complex transfer function and signal filtering across a broad range of total input AUCs. We further showed that both the qualitative nature of the transfer function and the quantitative maximum information content of the gene could be dramatically tuned by constitutive recruitment of chromatin regulators to the promoter. This work directly demonstrates the deep signal processing potential of a single individual gene and develops molecular and computational tools that can be used to harness it.

Epigenome editors, chromatin regulators fused to DNA-binding domains, are an increasingly important tool in both biological research and therapeutic development (Adamson et al., 2016; Keung et al., 2014; Liu et al., 2018; Park et al., 2019; Thakore et al., 2016). Their functions have been largely viewed as inducing static changes in state, for example in which alteration of histone modifications or recruitment of a transactivator/repressor might lead to up- or down-regulation of transcription. However, it is now clear that both the dynamic recruitment of editors themselves as well as their impact on the interpretation and processing of other dynamic signals can have profound regulatory effects, including the filtering of different types of dynamic signals well beyond just monotonic on or off control. Such properties have previously been shown to be tunable through mutations in proteins or alterations of protein scaffolds (Bashor et al., 2019; Hao et al., 2013). It is now evident that altering the epigenome can also regulate filtering properties in a potentially reversible way without changing gene or protein sequences. This could be used to confer useful functions such as expressing therapeutic proteins only within a specific range of input signals.

It is also clear that while the expression strength of an output signal can be tuned by altering the concentration of an input epigenome editor or TF using conventional inducible systems (i.e. LacI or TetR), this type of amplitude-based control may not always be ideal. For example, we found frequency modulation was able to confer a similar output dynamic range as amplitude modulation but with tighter population distributions and therefore greater mutual information and reliability. Furthermore, when combining all three dynamic parameters, mutual information was further increased, enhancing the amount of information that could reliably be transmitted by the gene. By achieving more possible output states for a limited number of inputs, tighter control over output responses is possible and may be particularly important in applications that are sensitive to expression levels such as regulating immune responses.

In addition to informing the design of synthetic biological tools like epigenome editors, this work suggests consideration of how both the fidelity and inherent transfer functions of natural signaling systems may exhibit considerable differences between cell types and/or over time. The transfer functions and the mutual information of the same individual genes may switch how they interpret dynamic signals in distinct cell types or in distinct cell states—or during the progression of cancer, aging, or normal development. Many natural systems shown to interpret dynamic signals may also alter their interpretations or transfer functions depending on time and space, including neural cell fate decision making (Imayoshi et al., 2013; Marshall, 1995) and cancer proliferation (Bugaj et al., 2018). Many other biological processes have been linked to dynamic pulsing, such as B-cell activation (Inoue et al., 2016) and responses to radiation (Purvis et al., 2012).

The exploration of dynamic signaling provides opportunities to continue shifting biological engineering to quantitative frameworks borrowed from disciplines in the physical sciences and engineering, but it also contributes new insights for those frameworks due to the distinctive properties of biological systems. For example, this work presents analogies to the concept of dynamic transfer functions common in process control theory, which formalizes the description and prediction of how outputs are controlled by input signals; yet, as we showed, a gene regulated by chromatin is a highly complex ‘unit process’ that can intriguingly morph its transfer function to have distinct filtering properties, without a change in gene sequence. Something analogous in a conventional unit process like a chemical reactor might, in contrast to a biological system, require drastic actions like altering the reactor’s material properties or shape.

Information theory also provides a theoretical basis to move from phenomenological frameworks of dose-dependent gene responses that assumes continuous and graded control over gene expression levels, to thinking about true information transmission more rigorously. Interestingly, we and others (Billing et al., 2019; Cheong et al., 2011; Dubuis et al., 2013; Grabowski et al., 2019; Hansen and O’Shea, 2015; Harton and Batchelor, 2017; Jetka et al., 2019; Selimkhanov et al., 2014; Tkacik et al., 2009; Tudelska et al., 2017; Uda et al., 2013) have shown that these biological unit processes have relatively low information content of less than 1.5 bits (i.e. ∼3 states). This may initially present a conundrum for how biological systems can exert such high-level functions within highly variable and complex environments. However, each gene can respond to multiple TFs and other factors including nucleosome remodelers while three-dimensional topology also impacts gene expression. Many promoters especially in mammalian systems can be quite large, promoting the ability to sense additional inputs. The diversity of multiple inputs could further increase the MI of genes. Furthermore, linking multiple components into higher order circuits can yield overall greater information transmission as well as lend precision or robustness to input-output responses (Barkai and Leibler, 1997). For all of these reasons, the MI of biological systems may be much higher than currently measured. As a case in point, the simple addition of just one additional input factor, recruiting CRs such as Arp8p or Rxt3p, was able to increase the MI of the reporter in our system (Figure 6). The ability to increase MI could lead to more complex biological sensors, while reducing MI could provide expression systems that are more robust to environmental stressors (Billing et al., 2019).

There are many exciting avenues to expand into and explore. In our work, to be able to screen large numbers of input signal patterns as well as CRs, we relied on endpoint measurements that could be rapidly measured by flow cytometry. However, information can also be stored in the dynamics of the output signal, e.g. the production rate, time-delay of repression/activation, or oscillatory behavior. High throughput approaches that can track the output dynamics of thousands of cultures would unlock this potential space for investigation. We also investigated a single promoter while different promoter structures would likely confer distinct transfer functions (Hansen and O’Shea, 2016). Additional factors that could be explored include the effect of gene duplications, tuning the binding kinetics and/or cooperativity of TFs, assessing species differences, and exploring the contribution of multiple inputs which would already have nice quantitative frameworks to build upon from process control theory (i.e. multiple input multiple output or “MIMO” control). Continued advances in experimental and computational systems that can handle the large parameter space of dynamic signals will unlock our ability to measure, quantify, and understand information transmission in biological systems, and reveal the underpinnings of how limited numbers of components can give rise to the rich complexity of biological functions.

## Acknowledgements

This work was supported by the NSF Emerging Frontiers in Research and Innovation program, the NCSU Provost’s Professional Experience Program (to NL), the NCSU Office of Undergraduate Research (to JYL and LC), Post 9/11 GI Bill (to JBL) and an NIH T32 Molecular Biotechnology Training Program Fellowship (to JBL and LC). We thank Mike Mantini for help in designing and building light matrices.

## STAR methods

### Resource Availability

#### Lead contact

Further information and requests for resources and reagents should be directed to and will be fulfilled by the Lead Contact, Albert J. Keung.

#### Materials availability

Plasmids and plasmid maps generated in this study will be deposited to Addgene upon publication.

#### Data and code availability

Data and code are available at github.com/keung-lab/Dynamic-Transfer-Functions.

### Experimental model and subject details

#### Cell culture

The background cell line for all experiments in this study was YPH500 *(α, ura3-52, lys2-801, ade2-101, trp1Δ63, his3Δ200, leu2Δ1)* (Stratagene). Cells were cultured in synthetic drop-out media or complete media made (Sunrise Scientific) with YN-B from Sigma and 2% w/v glucose. Our host strain was generated by genomically integrating an expression cassette that constitutively expresses TetR, LacI, and GEV (Louvion et al., 1993)(cloned into single-integrating plasmid pNH607 [*HO*]). Constitutive expression of the repressors in glucose-containing media ensured low basal levels of expression of ZF-CRY2 and CIB1-VP16 from the engineered *GAL1* promoters, which was relieved by the respective addition of the chemical inputs, ATc and IPTG, along with β-estradiol to the medium.

### Method details

#### Cloning and plasmid construction

All plasmid constructs were created using standard molecular biology techniques and Gibson isothermal assembly. Plasmids were grown and prepared from either NEB Turbo or Stable competent cells. The CR plasmid library was synthesized as previously described (Keung et al., 2014). In short, primer sequences were obtained from the *Saccharomyces* Genome Database (SGD). These primers (synthesized by Integrated DNA Technologies) were used to amplify full length CR ORFs from wild-type yeast (BY4742). SbfI and NotI flanking restriction sites were used to ligate the PCR products to the C-terminus of (3xFLAG)-(nuclear localization sequence)-(97-4 zinc finger array)-(17 amino acid glycine-serine linker) using plasmid pJL50.

#### Cell strain generation

Strains were constructed by sequential plasmid transformations using standard lithium acetate-based transformation techniques. Plasmids were first linearized using PmeI or SbfI. Following transformation, cells were grown on selective auxotrophic minimal media (Sunrise). Strains are listed in Table 1 while plasmids are listed in Table 2.

**Table 1.** Yeast strains and integrating plasmid constructs.

|  | Marker loci |  |  |  |  |  |
| --- | --- | --- | --- | --- | --- | --- |
| Strain ID | *HO* | *URA3* | *TRP4* * | *LEU2* | *HIS3* | Figure |
| Y11 | pNH607 |  |  |  |  | 1B, 2, 3, 4, 5, 6 |
| JY28 | pNH607 |  |  | pJL29 |  |  |
| JY29 | pNH607 | pJL30 |  | pJL29 |  | 1B |
| JY138 | pNH607 | pJL30 |  | pJL29 | pNL2 | 1B |
| JY145 | pNH607 | pJL30 | pJL38 | pJL29 | pJL32 | 1B, 2, 3, 4, 5, 6 |
| CR library | pNH607 | pJL30 | pJL50-EE | pJL29 | pJL32 | 5, 6 |
| JY30 | pNH607 | pJL30 |  | pJL29 | pJL32 |  |

\* Trp4 auxotrophic marker constructs were integrated into AmpR of the *LEU2* plasmid.

**Table 2.** Plasmids.

| **Plasmid ID** | **Parent** | **Features** | **Method of construction** |
| --- | --- | --- | --- |
| pJL29 | pYL-78, pYL-26 | 2x43-8 ZFO-1x97-4 ZFO<br>pCyc mCherry tCyc | Cut both with ClaI and EagI. Ligate. |
| pJL30 | C28, pJL15 | pGal1-TetO-3xNLS-43-8ZF-Cry2-tADH1, URA3 selection<br>marker | PCR pJL15 with JLp161/162. Cut<br>PCR and pC28 with SpeI/MluI.<br>Ligate. |
| pJL38 | C1, pYL-26,<br>pYG-8 | 2x 43-8 ZFO 1x97-4 ZFO-pCyc1-mCitrine-tCyc | Cut C1 with KpnI/NotI. PCR pYL-26<br>with JLp251/JYLp4. PCR pJL31 with<br>JYLp5/JYLp6. Gibson assembly. |
| pJL32 | C6, pJL1 | pGal1-LacO-3NLS-CIB1-VP16-tADH1 | PCR C6 with JLp163/JLp169. PCR<br>pJL1 with JLp165/166. PCR pJL1<br>with JLp167/JLp168. Gibson<br>assembly |
| pNL2 | pJL32 | pGal1-LacO-3NLS-CIB1-tADH1 | PCR C6 with JLp163/JLp169. PCR<br>pJL1 with JLp165/JLp166. PCR pJL1<br>with JLp167/168. Gibson assembly. |
| pJL50-EE | C1, pAG18 | pTEF1-3NLS-97-4ZF-Sbfl-EE-NotI | PCR pAG18 with JLp342/JL343. Cut<br>C1 with KpnI/ApaI. Gibson<br>assembly. |
| pJL15 | C6, pGal4BD-CRY2 | pGal1-TetO-3xNLS-43-8ZF-Cry2-tADH1, HIS3 selection<br>marker | PCR pLi-2 with JLp46/47. Cut PCR<br>and C6 with Sbfl/NotI. Ligate with T4<br>ligase. |
| pJL31 | C1, pYL-26, pYG-8 ,+[pYL-26 JYLp7/8] | pCyc1-mCitrine-tCyc1 | Cut C1 with KpnI/NotI. PCR pYL-26 with JYLp3/JYLp4 and JYLp7/JYLp8. PCR pYG-8 with JYLp5/JYLp6. Gibson assembly. |
| pJL1 | C8, pGal4AD-CIB1 | pGal1-LacO-3NLS-VP16-CIB1-tADH1 | PCR pLi-1 with JLP1/2. Cut PCR and C8 with XmaI/BamHI. Ligate. |
| pYL-78 |  | 2x 43-8 ZFO 1x97-4 ZFO-pCyc1-yEGFP-tCyc1, LEU2 selection marker |  |
| pYL-26 |  | 2x 43-8 ZFO 1x97-4 ZFO-pCyc1-ymCherry-tCyc1, URA3 selection marker |  |
| C28 |  | pGal1-LacO-3xNLS- 97-4ZF-VP16-tAdh1, URA3 selection marker |  |
| C1 |  | pGal1-LacO-3xNLS- 97-4AF-VP15-tADH1, TRP1 selection marker |  |
| pYG-8 |  | Yeast mCitrine |  |
| C6 |  | pGal1-tetO-3NLS- 43-8ZF-VP16-tADH1, HIS3 selection marker |  |
| pAG18 |  | pTef1-spSwi6-tTEF1 |  |
| C8 |  | pGal1-tetO-3NLS- 43-8ZF-VP15-tADH1, HIS3 selection marker |  |

pGal4BD-CRY2 and pGal4AD-CIB1 were gifts from Chandra Tucker. Addgene plasmid # 28243; http://n2t.net/addgene:28243; RRID:Addgene\_28243 (Kennedy et al., 2010).

**Table 3.**
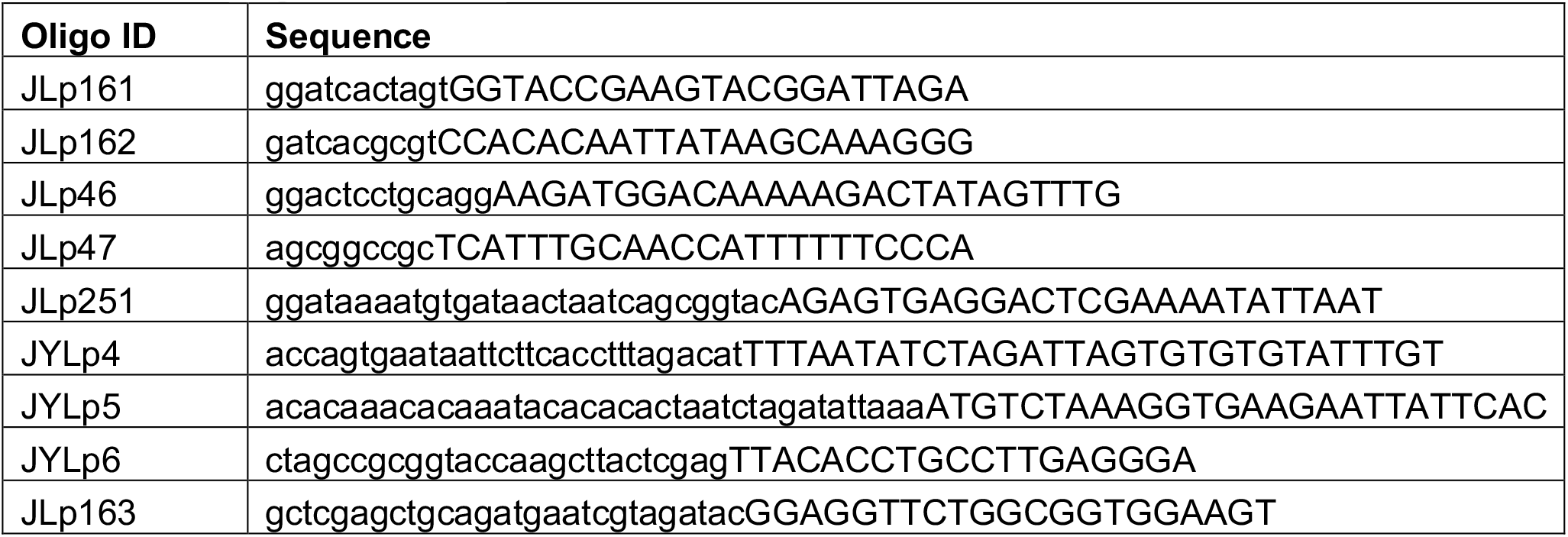
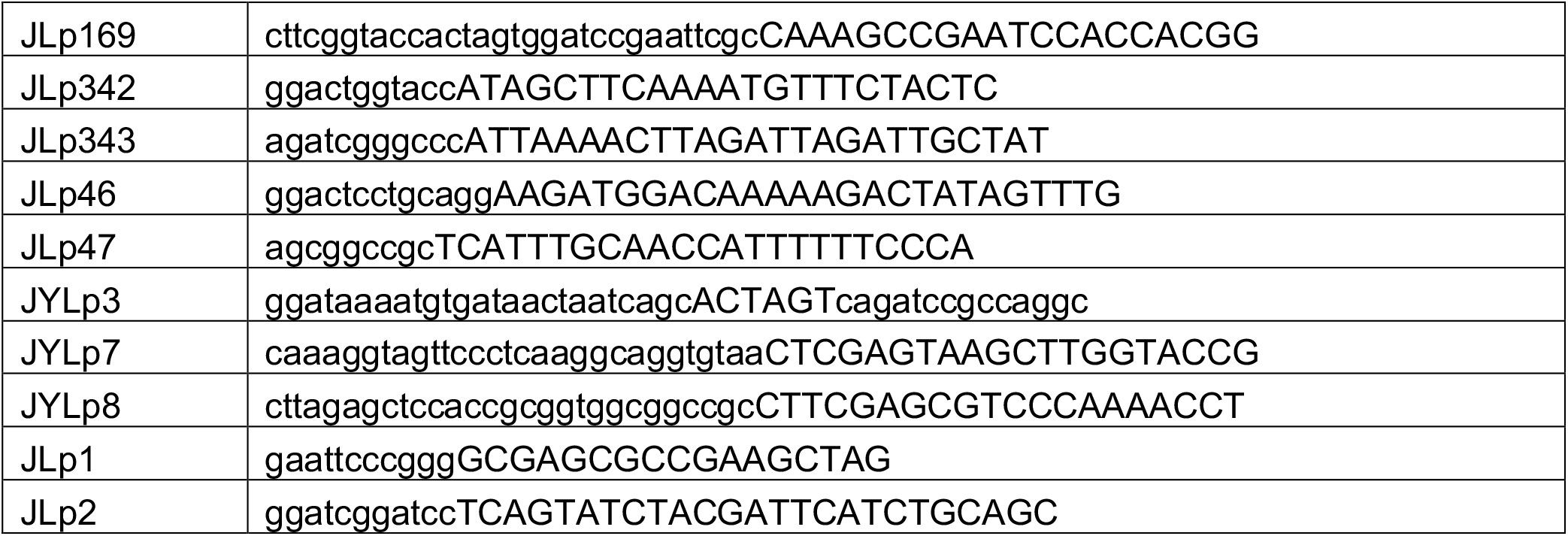
DNA oligomers.

#### LED matrix construction and calibration

Three LED matrices were made. Each had a LED housing unit 3D printed using black polylactic acid plastic. Each housing unit was designed to fit a standard 96-well plate with a single, programmable LED for each well. The plans for the housing unit were created in TinkerCad and are available upon request. Female socket pins were glued to the housing unit to connect to each LED. 60 or 92 blue LEDs (Chanzon, 100F5T-YT-WH-BL) were connected to 220 Ω resistors before being connected to 16-channel servo driver breakout boards (PCA9685, Adafruit). Three or five breakout boards were used for each 60 LED or 92 LED matrix, respectively. In addition, 12 LEDs were controlled directly from the PWM pins (0-11) on the Arduino Due. Each matrix was controlled by an Arduino Due, using I2C. Arduino code was written using the Arduino IDE to control the pulse width, intensity, and frequency of light pulses.

Calibration of the LEDs was done by attaching each LED matrix to a black 96-well plate with a flat, clear bottom (Corning, 3788) and taking 59 images across each well using a microscope (Nikon Ti-Eclipse, 20x SP objective, z=4486 µm) through a DAPI filter cube (Chroma Technology, 96360) with exposure time set to 100ms. The pixel intensity was extracted using a custom Matlab code. For a single well, pixel intensities for each image were read using the *imread* function. The total pixel intensities for each image were summed and then divided by the number of images. The average intensity was also determined for a well without an LED. This value was subtracted from all wells’ intensities to get the working LED intensity. This was done for three Arduino inputs and fit to a line for each well. The calculated values were used as initial inputs for the intensities used for the experiments. The intensities were then checked and adjusted before each experiment to be within 20 percent of the desired intensity.

#### LED intensity measurement with power meter

The LED intensities can be converted to nW using Figure S1 panel G. The power meter measurements were taken using a PM100D power meter (ThorLabs) with a S140C probe. A M134L01 fiber patch cable (0600 µm core, 0.5 NA, FC/PC to SMA, 1 m length) was connected to the probe via the FC/PC connector. For each well, the SMA connector was held against the bottom of a clear, flat-bottom plate (Corning, 3788) connected to the LED matrix. Multiple readings were taken at various locations for each well, and the mean was plotted and fitted to a line as shown in Figure S1 panel G.

#### Flow cytometry

Yeast colonies were picked from plates and cultured 24-48 h in the appropriate auxotrophic SD media. Cultures were diluted to ∼0.1 OD600 with auxotrophic dropout media that contained 0.4 µg/mL ATc, 10mM IPTG, 5µM of beta-estradiol, and 0.02 mg/mL adenine. Cells were incubated at 30°C and 900 RPM, in the dark, for 8-9 h to allow for expression of ZF-CRY2 and CIB1-VP16. Cells were then diluted 1:30 with 200 µL SD-complete media, containing the same chemicals as above, into U-bottom, black 96-well plates (Costar, 3792). Samples were prepared as much as possible in a red light environment to reduce premature binding of CIB1 and CRY2. Plates were attached to the LED matrices and incubated at 30°C for 14 h at 500 RPM. Replicate plates were grown in the dark. The shaking speed was reduced to prevent damage of the LED matrices.

Prior to flow cytometry, 100 µL of 0.03 mg/mL of cycloheximide was added to each sample. Samples were then incubated in the dark at room temperature for 1 h to allow for mCherry maturation. Fluorescent measurements were taken using a MACSQuant VYB (Miltenyi Biotec). A maximum of 20,000 events were collected per sample. Plates were stored at 4°C while waiting for other plates to be run on flow cytometer. All samples were run within 8 h of adding cycloheximide.

All samples were gated using SSC-A and FSC-A, using a custom MATLAB code based on methods described previously (Newman et al., 2006). To summarize, the FSC-A and SSC-A were natural log transformed. Cell outside a circle of radius 0.7 around the median FSC-A and SSC-A were excluded from further analyses. Any gated samples with less than 250 events were also excluded from further analyses.

### Quantification and statistical analysis

#### Fold change and noise calculation

The population medians of the fluorescence distributions were calculated for the gated populations. The autofluorescence value of *S. cerevisiae* YPH500 cells harboring no genomic integrations was subtracted from these values. “Fold change” values were calculated as the ratio of fluorescence values from cells exposed to a given blue light pattern to those from cells grown without blue light. Four isogenic strains were grown for each light condition. The “coefficient of variation”, or CV is the robust CV calculated using the equation: 0.5 * [intensity(at 84.13 percentile) - intensity(at 15.87 percentile)] / median. Outliers were identified using MATLAB’s isoutlier function, which classifies values as an outlier if it is more than three scaled median absolute deviations away from the median fold change or CV. Any outliers were excluded from the means graphed in Figures 2-3.

To minimize the variability due to the large number of plates in the CR screens (Figure 6), each plate with blue light was normalized to the strain with VP16 only (JY145) with light always on and light intensity at 6×10^10^ au, which was grown in the same plate. Each plate without blue light was normalized to VP16 only (JY145) with no light, grown in the same plate. Population medians were used to calculate the fold change, which is the fluorescence of the strain with light divided by fluorescence of strain without light.

#### Maximal mutual information calculation

The maximal mutual information was found as previously described in (Cheong et al., 2011; Hansen and O’Shea, 2015; Shannon, 1948). For each sample, events were gated as described in the *Flow Cytometry* section. Then the mCherry measurements were normalized to the FSC-A measurements. The responses were discretized using logarithmically sized bins. The mutual information I(R;S), measured in bits, was calculated by

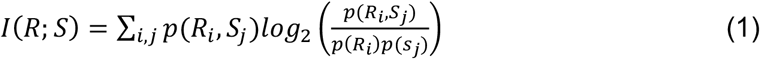

where S is the signal input and R is the observed response output. The response distribution is given by p(R), and p(s) is unknown. The maximal mutual information was found by solving the optimization in Equation 2.

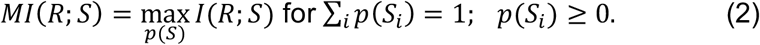

The above optimization was solved using the Blahut-Arimoto algorithm from code written by Piyush Singh (Singh, 2015). The MI was corrected for bias due to the number of bins by varying the number of bins from 5 to 50. The unbiased MI was calculated as the mean of MIs calculated using 21-41 bins, which is within the plateau region of MI versus number of bins. The MI was then corrected for under-sampling using jackknife sampling as described previously (Cheong et al., 2011; Hansen and O’Shea, 2015; Slonim et al., 2005). The means shown in Figures 5 and 6 are of the unbiased MIs from 3-4 isogenic strains.

#### Determining signal filtering clusters

Clustering was completed on individual replicates for all the epigenome editors. To discover the clusters depicted in Figure 6, strains that had low fold change and low variability of fold change among frequencies were removed. The fold changes were then logarithmically transformed. The remaining strains were grouped into 5 clusters using the *kmeans* function in MATLAB with correlation as the distance metric. The centroids from the resulting clusters were slightly modified to fit the behaviors in Figure 6C. The centroids are as follows: Cluster 1: 0.756341421875678, - 0.3, -0.338099818683020; Cluster 2: -0.736510820962514, 0.102405013200597, 0.65; Cluster 3: -0.351484685217366, 0.769986528990965, -0.418501843773599; Cluster 4: - 0.775512363928304, 0.550835601762814,0.3; Cluster 5: 0.0178768566235528, - 0.685625005390339, 0.667748148766786. The fold changes were then reclustered using these centroids with the *pdist2* function, again with correlation as the distance metric.

#### Statistical analyses

N-way ANOVA tests were performed using the *nanova* function in MATLAB. For the comparison among multiple conditions, a Tukey’s honest significant difference criterion (T-K analysis) was used via the *multcompare* function in MATLAB with a 95 percent confidence interval. The analysis of covariance (ANCOVA) was performed using the *aoctool* function.

#### Model Selection

Seventeen different three state and four state models were tested with a variety of architectures and between 5 and 9 fitted parameters. The models were validated using two metrics: first, the residual sum of the squares on the model outputted endpoints and experimental endpoints; and second, cross validation to the expected time course curve shape based on literature (Hansen & O’Shea 2013, Harton 2019, Benzinger & Khammash 2018). We found that a prior three-state model did not provide a sufficiently smooth output curve as compared to a four-state model (Figure S3B). We also varied the number and placement of Hill functions between models and found that only using a Hill function to describe the transition between P_ubound_ and P_bound_ was able to best replicate the endpoint behavior seen at low PWs (Figure S3C).

#### Deterministic Model Construction

To better understand the relationship between dynamic inputs and gene expression outputs in our system, a deterministic kinetic model was created, which is described by the following set of ODEs:

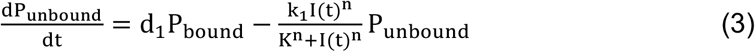

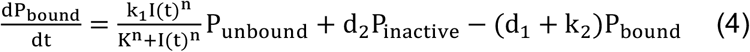

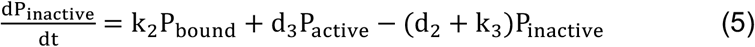

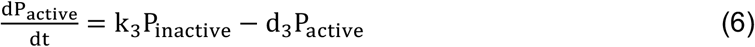

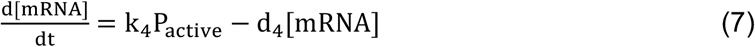

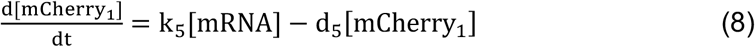

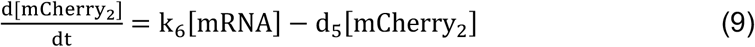

Here, d_1_ and k_1_ are the transition rates between the unbound and bound states, d_2_ and k_2_ are the transition rates between the bound and inactive states, and d_3_ and k_3_ are the transition rates between the inactive and active states. The transcription, translation, and maturation rates are k_4_, k_5_, and k_6_, respectively. The mRNA and mCherry degradation rates are d_4_ and d_5_. 8 were experimental constants, d_1_, k_1_, n, K, d_4_, k_5_, d_5_, and k_6_, and 5 were model fitted parameters, d_2_, k_2_, d_3_, k_3_, and k_4_. The fit of the model was assessed using the coefficient of determination. P_unbound_, P_bound_, P_active_, and P_inactive_ represent the probability of the promoter being in a given state and are each between 0 and 1 and must sum to 1 at any point in time. A Hill function was used to describe the transition between P_unbound_ and P_bound_. The input function is I(t), and is based on the PW, A, and F of the light condition. The light inputs were smoothed to prevent discontinuities and undefined values in the code, and the input amplitudes were 6e1 to 6e2 au rather than 6e9 to 6e10 au to prevent overflow error. Periods of the light being on were described by A(1 −e^− ct^), where A is the desired amplitude of the light condition and c = 0.6, for the entire pulse width (Hansen & O’Shea 2013). At the end of the pulse, the light was smoothed off using Be^− ct^, where B is the maximum amplitude reached during the pulse and c = 0.6, for the entire duration of the dark period (Hansen & O’Shea 2013).

#### Parameter Screen and Model Fitting

To fit the model to the data, sets of parameters fit by the model (d_2_, k_2_, d_3_, k_3_, and k_4_) were stochastically generated using Latin Hypercube Sampling (LHS). The ODEs were solved numerically using odeint in Python, and the model output at 14 hours was compared to the experimental value using the residual sum of the squares. The fitting was performed in two steps: initially, 1000 randomly generated sets of parameters, each sampled over a range of 0.0002 – 0.02 sec^-1^, was run through the model. Then, the parameter set that resulted in the highest R^2^ value was used to “fine-tune” the LHS sampling range, and 100 new sets were generated and run through the model. The fine-tuning processes was repeated four times, resulting in a R^2^ of 0.835 (Figure 3B). The model fitted parameters are shown in Figure S3A.

The experimental parameters unique to mCherry (d_4_, k_5_, d_5_, and k_6_), were found by randomly sampling within ranges provided by literature and fit to the 5 sec, 120 sec, and 600 sec PW experimental data while all other parameters were held constant. 1000 parameter sets were tested, with d_4_ ranging from 0.0012 – 0.023 sec^-1^, k_5_ from 0.2 – 0.3 sec^-1^, d_5_ from 0.000017 – 0.0013 sec^-1^, and k_6_ from 0.00084 – 0.002 sec^-1^ (Hansen & O’Shea 2013), and the best parameter set was chosen by comparing model to experimental endpoints. Experimental parameters unique to the blue light optogenetic system (d_1_, k_1_, n, and K) were fit based around the literature ranges of the dissociation and association rates of the system. d_1_ was found by sampling within 0.003 - 0.004 sec^-1^ and comparing the model and experimental endpoints. 11,000 parameter sets were generated with d_1_ and other model fitted parameters changing, and the parameter set that resulted in the highest R^2^ value for the entire data set was used to find the value for d_1_, which remained fixed for all fine-tuning parameter sets. The total forward on rate between and P_unbound_ and P_bound_ has been reported around 0.054 – 0.11 sec^-1^, K between 100 – 2500 and n between 0.5 – 4 (Rademacher 2017, Hansen & O’Shea 2013, Gonze & Abou-Jaoude 2013). k_1_ was fixed at 1.184 sec-1, K at 1400, and n at 1.5 so that the total forward rate was 0.01 sec^-1^ at I(t) = 60 au and 0.26 sec^-1^ at I(t) = 600 au. All model fitted parameters used in Figure 3B, as well as the experimental constants, are shown in Figure S3A.

## KEY RESOURCES TABLE

| REAGENT or RESOURCE | SOURCE | IDENTIFIER |
| --- | --- | --- |
| Antibodies |  |  |
| None |  |  |
| Bacterial and Virus Strains |  |  |
| NEB Turbos |  |  |
| NEB Stables |  |  |
| Biological Samples |  |  |
| None |  |  |
| Chemicals, Peptides, and Recombinant Proteins |  |  |
| Anhydrotetracycline | Fisher Scientific | Cat# 50-595-757 |
| IPTG | Fisher Scientific | Cat# BP175510 |
| Beta-estradiol | Fisher Scientific | Cat#AAL0380103 |
| Cycloheximide | Fisher Scientific | Cat#AAJ6690103 |
| YNB+nitrogen | Sigma | Cat# Y0626 |
| CSM | Sunrise Scientific | Cat#1001-100 |
| CSM-LEU | Sunrise Scientific | Cat#1005-010 |
| CSM-LEU-URA | Sunrise Scientific | Cat#1038-010 |
| CSM-LEU-HIS | Sunrise Scientific | Cat#1011-010 |
| CSM-LEU-HIS-URA | Sunrise Scientific | Cat#1015-010 |
| CSM-LEU-HIS-URA-TRP | Sunrise Scientific | Cat#1002-010 |
| Adenine hemisulfate | Sunrise Scientific | Cat#1905-010 |
| Critical Commercial Assays |  |  |
| None |  |  |
| Deposited Data |  |  |
| Github.com/keung-lab/Dynamic-Transfer-Functions |  |  |
| Experimental Models: Cell Lines |  |  |
| None |  |  |
| Experimental Models: Organisms/Strains |  |  |
| YPH500 | Stratagene |  |
| Oligonucleotides |  |  |
| See Oligonucleotides table |  |  |
| Recombinant DNA |  |  |
| See Plasmids table |  |  |
| Software and Algorithms |  |  |
| MATLAB 2018b | MathWorks | https://www.mathworks.com/downloads/ |
| Arduino IDE | Arduino | https://www.arduino.cc/en/Main/Software |
| TinkerCAD | Autodesk | https://www.tinkercad.com/ |
| Other |  |  |

## References

Adamson, B., Norman, T.M., Jost, M., Cho, M.Y., Nunez, J.K., Chen, Y.W., Villalta, J.E., Gilbert, L.A., Horlbeck, M.A., Hein, M.Y., et al. (2016). A Multiplexed Single-Cell CRISPR Screening Platform Enables Systematic Dissection of the Unfolded Protein Response. Cell 167, 1867-+.

An-adirekkun, J., Stewart, C.J., Geller, S.H., Patel, M.T., Melendez, J., Oakes, B.L., Noyes, M.B., and McClean, M.N. (2020). A yeast optogenetic toolkit (yOTK) for gene expression control in Saccharomyces cerevisiae. Biotechnology and Bioengineering 117, 886–893.

Bar-Even, A., Paulsson, J., Maheshri, N., Carmi, M., O’Shea, E., Pilpel, Y., and Barkai, N. (2006). Noise in protein expression scales with natural protein abundance. Nature Genetics 38, 636–643.

Barkai, N., and Leibler, S. (1997). Robustness in simple biochemical networks. Nature 387, 913–917.

Bashor, C.J., Patel, N., Choubey, S., Beyzavi, A., Kondev, J., Collins, J.J., and Khalil, A.S. (2019). Complex signal processing in synthetic gene circuits using cooperative regulatory assemblies. Science 364, 593-+.

Batchelor, E., Loewer, A., Mock, C., and Lahav, G. (2011). Stimulus-dependent dynamics of p53 in single cells. Molecular Systems Biology 7, 8.

Behar, M., and Hoffmann, A. (2010). Understanding the temporal codes of intra-cellular signals. Current Opinion in Genetics & Development 20, 684–693.

Benzinger, D., and Khammash, M. (2018). Pulsatile inputs achieve tunable attenuation of gene expression variability and graded multi-gene regulation. Nature Communications 9, 10.

Billing, U., Jetka, T., Nortmann, L., Wundrack, N., Komorowski, M., Waldherr, S., Schaper, F., and Dittrich, A. (2019). Robustness and Information Transfer within IL-6-induced JAK/STAT Signalling. Communications Biology 2.

Bintu, L., Yong, J., Antebi, Y.E., McCue, K., Kazuki, Y., Uno, N., Oshimura, M., and Elowitz, M.B. (2016). Dynamics of epigenetic regulation at the single-cell level. Science 351, 720–724.

Bugaj, L.J., Sabnis, A.J., Mitchell, A., Garbarino, J.E., Toettcher, J.E., Bivona, T.G., and Lim, W.A. (2018). Cancer mutations and targeted drugs can disrupt dynamic signal encoding by the Ras-Erk pathway. Science 361, 892-+.

Cai, L., Dalal, C.K., and Elowitz, M.B. (2008). Frequency-modulated nuclear localization bursts coordinate gene regulation. Nature 455, 485–U416.

Chen, S.Y., Osimiri, L.C., Chevalier, M., Bugaj, L.J., Nguyen, T.H., Greenstein, R.A., Ng, A.H., Stewart-Ornstein, J., Neves, L.T., and El-Samad, H. Optogenetic Control Reveals Differential Promoter Interpretation of Transcription Factor Nuclear Translocation Dynamics. Cell Systems. Chen, S.Y., Osimiri, L.C., Chevalier, M., Bugaj, L.J., Nguyen, T.H., Greenstein, R.A., Ng, A.H., Stewart-Ornstein, J., Neves, L.T., and El-Samad, H. (2020). Optogenetic Control Reveals Differential Promoter Interpretation of Transcription Factor Nuclear Translocation Dynamics. Cell Systems.

Cheong, R., Rhee, A., Wang, C.J., Nemenman, I., and Levchenko, A. (2011). Information Transduction Capacity of Noisy Biochemical Signaling Networks. Science 334, 354–358.

Dalal, C.K., Cai, L., Lin, Y.H., Rahbar, K., and Elowitz, M.B. (2014). Pulsatile Dynamics in the Yeast Proteome. Current Biology 24, 2189–2194.

Dubuis, J.O., Tkacik, G., Wieschaus, E.F., Gregor, T., and Bialek, W. (2013). Positional information, in bits. Proceedings of the National Academy of Sciences of the United States of America 110, 16301–16308.

Eldar, A., and Elowitz, M.B. (2010). Functional roles for noise in genetic circuits. Nature 467, 167–173.

Elowitz, M.B., and Leibler, S. (2000). A synthetic oscillatory network of transcriptional regulators. Nature 403, 335–338.

Grabowski, F., Czyz, P., Kochanczyk, M., and Lipniacki, T. (2019). Limits to the rate of information transmission through the MAPK pathway. Journal of the Royal Society Interface 16.

Gregor, T., Tank, D.W., Wieschaus, E.F., and Bialek, W. (2007). Probing the limits to positional information. Cell 130, 153–164.

Hansen, A.S., and O’Shea, E.K. (2013). Promoter decoding of transcription factor dynamics involves a trade-off between noise and control of gene expression. Molecular Systems Biology 9, 14.

Hansen, A.S., and O’Shea, E.K. (2015). Limits on information transduction through amplitude and frequency regulation of transcription factor activity. Elife 4.

Hansen, A.S., and O’Shea, E.K. (2016). Encoding four gene expression programs in the activation dynamics of a single transcription factor. Current Biology 26, R269–R271.

Hao, N., Budnik, B.A., Gunawardena, J., and O’Shea, E.K. (2013). Tunable Signal Processing Through Modular Control of Transcription Factor Translocation. Science 339, 460–464.

Hao, N., and O’Shea, E.K. (2012). Signal-dependent dynamics of transcription factor translocation controls gene expression. Nature Structural & Molecular Biology 19, 31–U47.

Harton, M.D., and Batchelor, E. (2017). Determining the Limitations and Benefits of Noise in Gene Regulation and Signal Transduction through Single Cell, Microscopy-Based Analysis. Journal of Molecular Biology 429, 1143–1154.

Harton, M.D., Koh, W.S., Bunker, A.D., Singh, A., and Batchelor, E. (2019). p53 pulse modulation differentially regulates target gene promoters to regulate cell fate decisions. Molecular Systems Biology 15, 15.

Imayoshi, I., Isomura, A., Harima, Y., Kawaguchi, K., Kori, H., Miyachi, H., Fujiwara, T., Ishidate, F., and Kageyama, R. (2013). Oscillatory Control of Factors Determining Multipotency and Fate in Mouse Neural Progenitors. Science 342, 1203–1208.

Inoue, K., Shinohara, H., Behar, M., Yumoto, N., Tanaka, G., Hoffmann, A., Aihara, K., and Okada-Hatakeyama, M. (2016). Oscillation dynamics underlie functional switching of NF-kappa B for B-cell activation. Npj Systems Biology and Applications 2, 9.

Jetka, T., Nienaltowski, K., Winarski, T., Blonski, S., and Komorowski, M. (2019). Information-theoretic analysis of multivariate single-cell signaling responses. Plos Computational Biology 15.

Kennedy, M.J., Hughes, R.M., Peteya, L.A., Schwartz, J.W., Ehlers, M.D., and Tucker, C.L. (2010). Rapid blue-light-mediated induction of protein interactions in living cells. Nature Methods 7, 973–U948.

Keung, A.J., Bashor, C.J., Kiriakov, S., Collins, J.J., and Khalil, A.S. (2014). Using Targeted Chromatin Regulators to Engineer Combinatorial and Spatial Transcriptional Regulation. Cell 158, 110–120.

Khalil, A.S., Lu, T.K., Bashor, C.J., Ramirez, C.L., Pyenson, N.C., Joung, J.K., and Collins, J.J. (2012). A Synthetic Biology Framework for Programming Eukaryotic Transcription Functions. Cell 150, 647–658.

Kouzarides, T. (2007). Chromatin modifications and their function. Cell 128, 693–705.

Lee, T.I., and Young, R.A. (2013). Transcriptional Regulation and Its Misregulation in Disease. Cell 152, 1237–1251.

Li, B., Carey, M., and Workman, J.L. (2007). The role of chromatin during transcription. Cell 128, 707–719.

Liu, H.T., Yu, X.H., Li, K.W., Klejnot, J., Yang, H.Y., Lisiero, D., and Lin, C.T. (2008). Photoexcited CRY2 Interacts with CIB1 to Regulate Transcription and Floral Initiation in Arabidopsis. Science 322, 1535–1539.

Liu, Y.X., Yu, C., Daley, T.P., Wang, F.Y., Cao, W.S., Bhate, S., Lin, X.Q., Still, C., Liu, H.L., Zhao, D.H., et al. (2018). CRISPR Activation Screens Systematically Identify Factors that Drive Neuronal Fate and Reprogramming. Cell Stem Cell 23, 758-+.

Louvion, J.F., Havauxcopf, B., and Picard, D. (1993). FUSION OF GAL4-VP16 TO A STEROID-BINDING DOMAIN PROVIDES A TOOL FOR GRATUITOUS INDUCTION OF GALACTOSE-RESPONSIVE GENES IN YEAST. Gene 131, 129–134.

Maheshri, N., and O’Shea, E.K. (2007). Living with noisy genes: How cells function reliably with inherent variability in gene expression. Annual Review of Biophysics and Biomolecular Structure 36, 413–434.

Marshall, C.J. (1995). SPECIFICITY OF RECEPTOR TYROSINE KINASE SIGNALING - TRANSIENT VERSUS SUSTAINED EXTRACELLULAR SIGNAL-REGULATED KINASE ACTIVATION. Cell 80, 179–185.

Newman, J.R.S., Ghaemmaghami, S., Ihmels, J., Breslow, D.K., Noble, M., DeRisi, J.L., and Weissman, J.S. (2006). Single-cell proteomic analysis of S-cerevisiae reveals the architecture of biological noise. Nature 441, 840–846.

Park, M., Patel, N., Keung, A.J., and Khalil, A.S. (2019). Engineering Epigenetic Regulation Using Synthetic Read-Write Modules. Cell 176, 227-+.

Polstein, L.R., and Gersbach, C.A. (2015). A light-inducible CRISPR-Cas9 system for control of endogenous gene activation. Nature Chemical Biology 11, 198–U179.

Purvis, J.E., Karhohs, K.W., Mock, C., Batchelor, E., Loewer, A., and Lahav, G. (2012). p53 Dynamics Control Cell Fate. Science 336, 1440–1444.

Purvis, J.E., and Lahav, G. (2013). Encoding and Decoding Cellular Information through Signaling Dynamics. Cell 152, 945–956.

Rademacher, A., Erdel, F., Trojanowski, J., Schumacher, S., and Rippe, K. (2017). Real-time observation of light-controlled transcription in living cells. Journal of Cell Science 130, 4213–4224.

Rosenfeld, N., Young, J.W., Alon, U., Swain, P.S., and Elowitz, M.B. (2005). Gene regulation at the single-cell level. Science 307, 1962–1965.

Schielzeth, H. (2010). Simple means to improve the interpretability of regression coefficients. Methods in Ecology and Evolution 1, 103–113.

Selimkhanov, J., Taylor, B., Yao, J., Pilko, A., Albeck, J., Hoffmann, A., Tsimring, L., and Wollman, R. (2014). Accurate information transmission through dynamic biochemical signaling networks. Science 346, 1370–1373.

Shannon, C.E. (1948). A MATHEMATICAL THEORY OF COMMUNICATION. Bell System Technical Journal 27, 623–656.

Singh, P. (2015). BlahutArimoto.m (GitHub).

Slonim, N., Atwal, G.S., Tkacik, G., and Bialek, W. (2005). Information-based clustering. Proceedings of the National Academy of Sciences of the United States of America 102, 18297–18302.

Thakore, P.I., Black, J.B., Hilton, I.B., and Gersbach, C.A. (2016). Editing the epigenome: technologies for programmable transcription and epigenetic modulation. Nature Methods 13, 127–137.

Tkacik, G., Walczak, A.M., and Bialek, W. (2009). Optimizing information flow in small genetic networks. Physical Review E 80.

Tudelska, K., Markiewicz, J., Kochanczyk, M., Czerkies, M., Prus, W., Korwek, Z., Abdi, A., Blonski, S., Kazmierczak, B., and Lipniacki, T. (2017). Information processing in the NF-kappa B pathway. Scientific Reports 7, 14.

Uda, S., Saito, T.H., Kudo, T., Kokaji, T., Tsuchiya, T., Kubota, H., Komori, Y., Ozaki, Y.-i., and Kuroda, S. (2013). Robustness and Compensation of Information Transmission of Signaling Pathways. Science 341, 558–561.

