## Supplemental figures for "Mapping the dynamic transfer functions of epigenome editing"

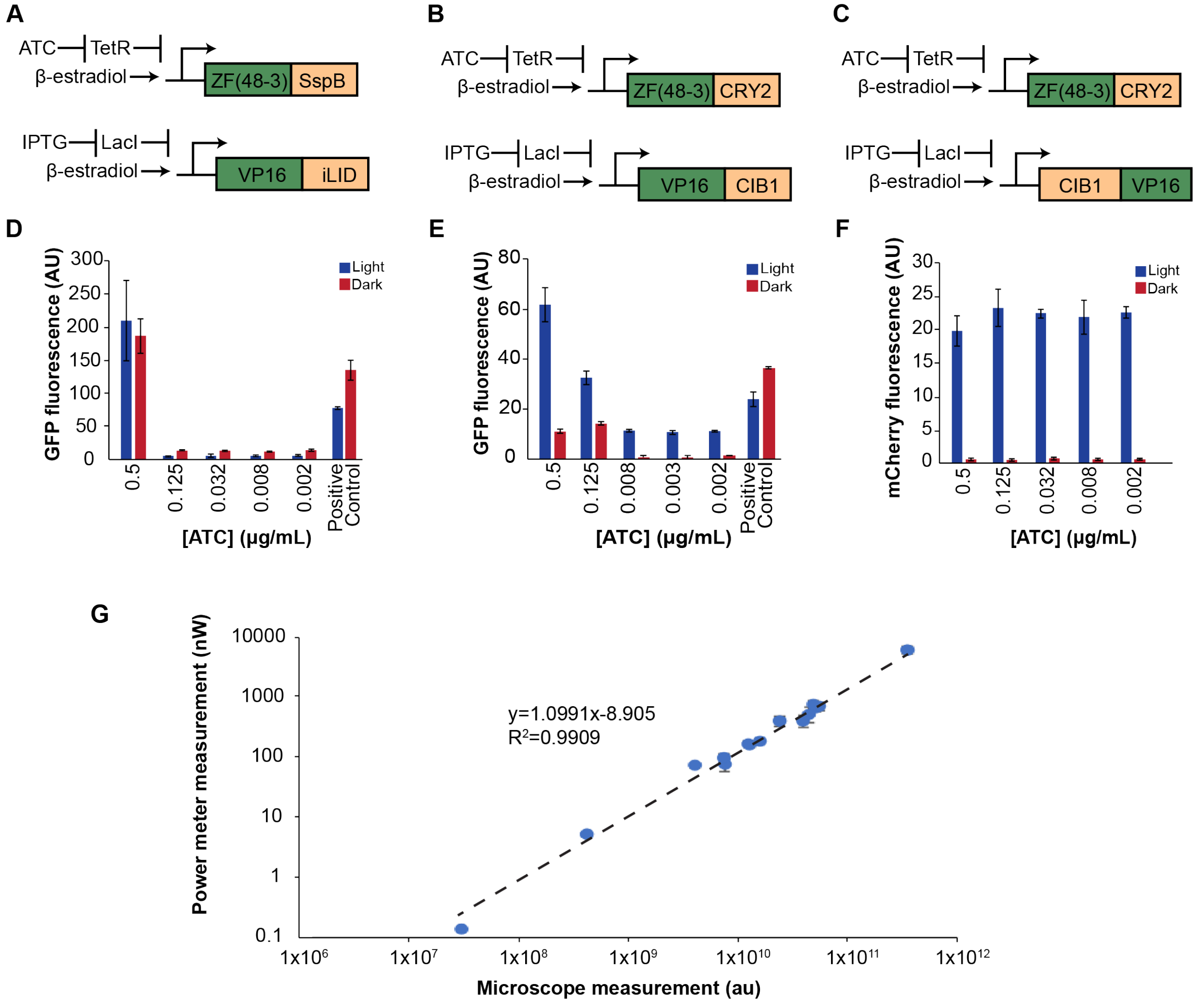


**Figure S1. Related to Figure 1. Development of optogenetic system.** (A-C) Schematics of DNA constructs tested. In addition to CRY2/CIB1, improved light-induced dimer (iLID) and its binding partner (SspB) were tested. (D) The iLID/SspB system showed no light-specific inducible activation. (E) Fluorescent output for ZF43-8-CRY2/VP16-CIB1 showed light-inducible activation. However, high ATC concentrations led to some activation without light. (F) Switching the fusion of CIB1 and VP16 produced robust light-inducible activation with minimal activation in the dark. The light condition was a single pulse (~6e10 au) for 6.5-7 hours. IPTG concentration was 20 mM for all plots. Error bars are 95% confidence intervals. Positive control was ZF43-8-VP16. (G) Light intensity relationship between microscope and power meter measurements provides a relationship between au and power. Equation on graph is a linear fit for $y=\log_{10}$(Power meter measurement) and $x=\log_{10}$(Microscope measurement).

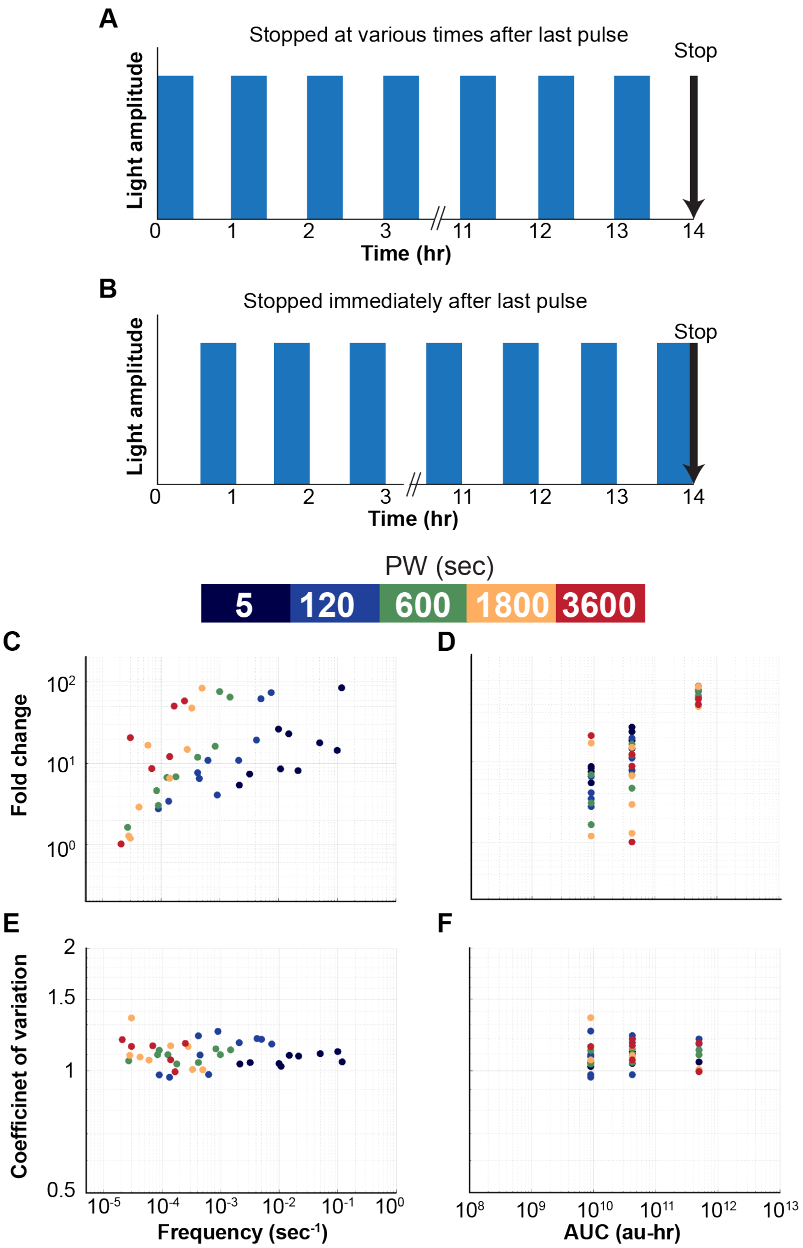


**Figure S2. Related to Figure 2 and Figure 4. Sampling time does not account for filtering.** To determine whether the time after the last light pulse affects the fold change, we tested two scenarios (A-B). (A) The light pulses began at the same time for all conditions, but the time after the last light pulse and addition of cycloheximide (CHX) varied according to equation: 1/F-PW. (B) The light pulses began at different times in order for them all to synchronously end at the same time. CHX was added to each sample immediately after the last pulse ended. (C-D) Results of the scenario shown in panel B, chosen for three AUCs. For results from panel A, see Figure 2. (C) Both scenarios exhibited linear trends of fold change versus F with higher PWs having higher slopes. (D) Importantly, the filtering behavior was still observed in scenario B where distinct mCherry outputs were achieved at the same AUCs. (E-F) Noise for scenario shown in panel B. Figure 5 shows the noise for panel A.

**A**

| Constant | Meaning | Value |
|---|---|---|
| $d_1$ | CRY2-CIB1 dissociation rate | $3.72\times10^{-3}$ sec$^{-1}$ |
| $k_1$ | CRY2-CIB1 binding rate constant | 1.184 sec$^{-1}$ |
| K | Activation Coefficient | 1400 |
| n | Hill Coefficient | 1.5 |
| $d_2$ | Inactive State to Bound State | $1.61\times10^{-2}$ sec$^{-1}$ |
| $k_2$ | Bound State to Inactive State | $3.46\times10^{-4}$ sec$^{-1}$ |

| Constant | Meaning | Value |
|---|---|---|
| $k_3$ | Inactive State to Active State | $1.42\times10^{-2}$ sec$^{-1}$ |
| $d_3$ | Active State to Inactive State | $1.69\times10^{-2}$ sec$^{-1}$ |
| $k_4$ | Transcription rate | $1.25\times10^{-2}$ sec$^{-1}$ |
| $d_4$ | mRNA degradation rate | $1.42\times10^{-2}$ sec$^{-1}$ |
| $k_5$ | Translation rate | $2.72\times10^{-1}$ sec$^{-1}$ |
| $d_5$ | mCherry degradation rate | $4.22\times10^{-5}$ sec$^{-1}$ |
| $k_6$ | mCherry maturation rate | $8.79\times10^{-4}$ sec$^{-1}$ |

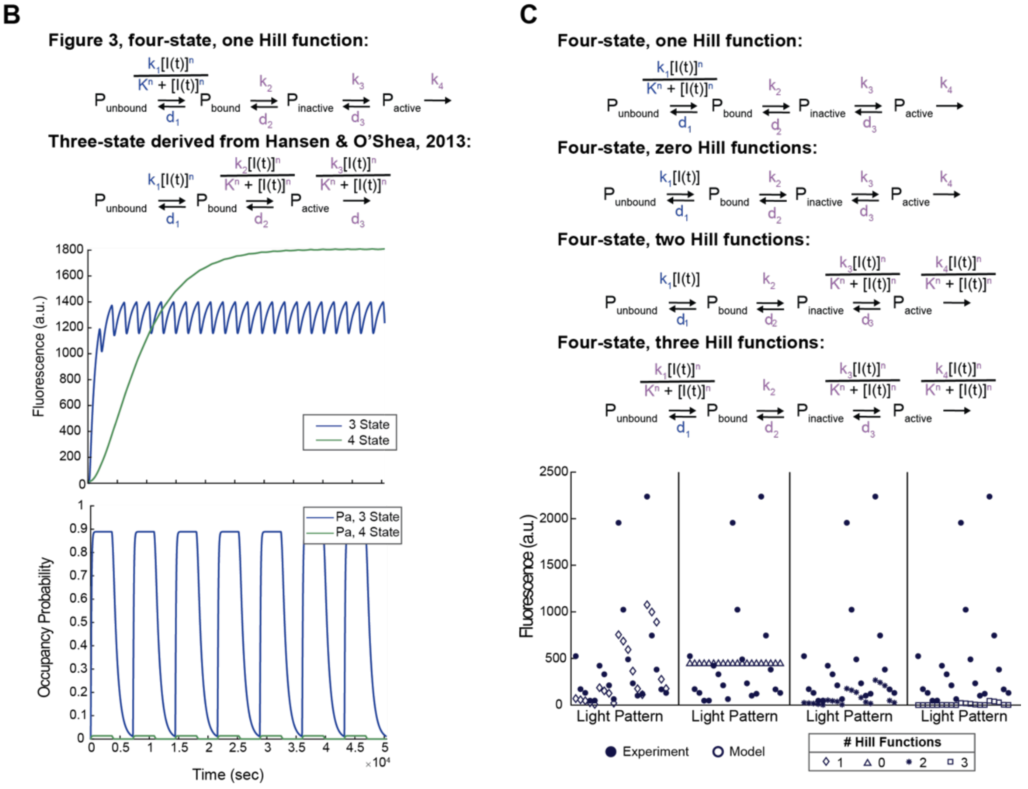


**Figure S3. Related to Figure 3. Comparison of different models.** (A) Values of fitted parameters for the four-state, one Hill function model. (B) Comparison of the four-state model used in Figure 3 and a previous three-state model. The three-state model exhibited significant oscillations as well as a very sharp initial increase in fluorescence that did not match experimental results. (C) Additional Hill functions were added to the model based upon strategies applied in prior work; however, the model with only one Hill function was able to best fit the experimental data.

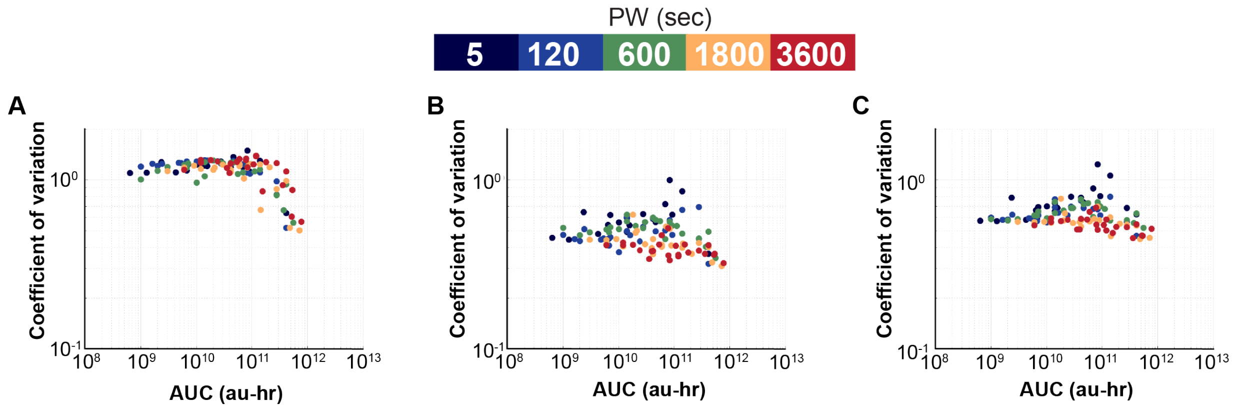


**Figure S4. Related to Figure 4. Effects of size gating and normalization on noise.** (A) Noise with FSC-A vs SSC-A gated with a large radius (0.7 with logarithmically transformed data). (B) Noise with small radius gate (0.3 with logarithmically transformed data). (C) Noise with large gating radius and fluorescence divided by FSC-A.

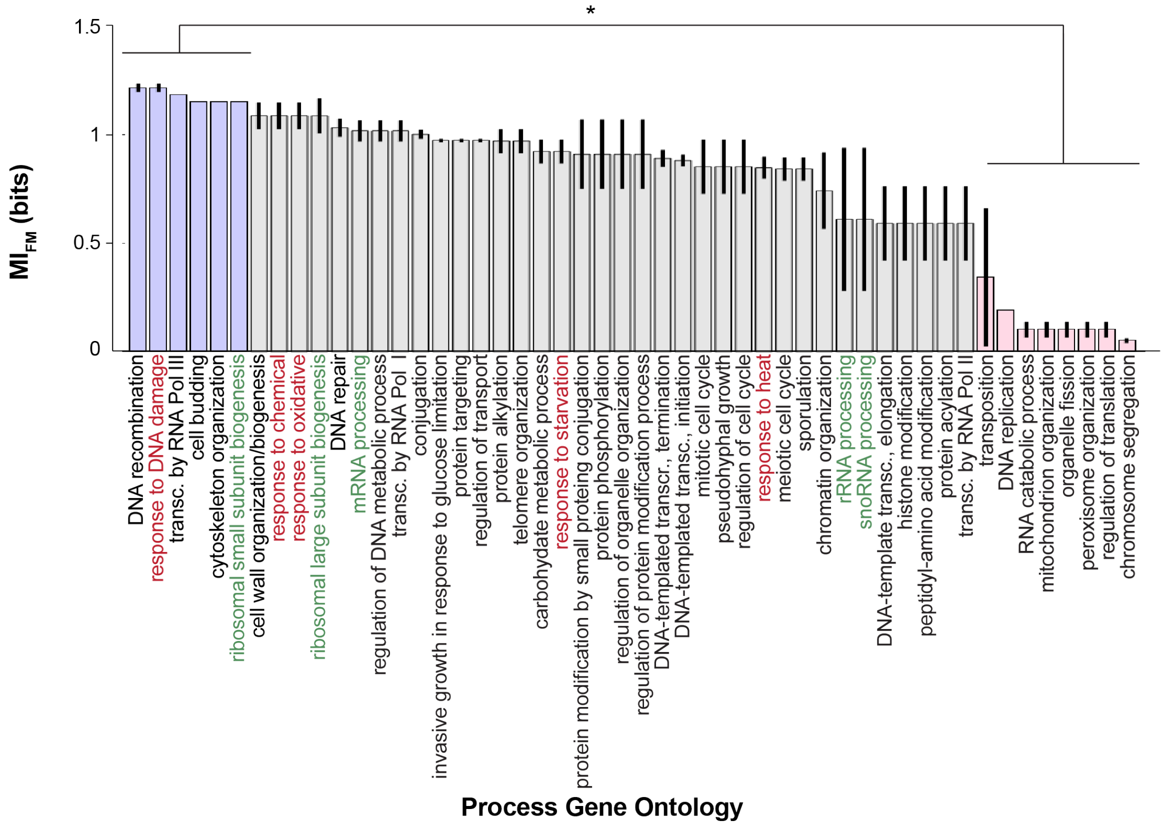


**Figure S5. Related to Figure 6. Gene ontology.** Average $MI_{FM}$ for each Process gene ontology term group. Chromatin regulators for each gene ontology were determined using the genes in the Yeast Genome Database (Cherry et al., 2012).

Cherry, J., Hong, E., Amundsen, C., Balakrishnan, R., Binkley, G., Chan, E., . . . Wong, E. (2012). Saccharomyces Genome Database: the genomics resource of budding yeast. Retrieved 20201209
